# Co-chaperone-mediated post-translational control of efflux pump induction underlies adaptive β-lactam resistance in *Caulobacter crescentus*

**DOI:** 10.1101/2023.02.20.529182

**Authors:** Jordan Costafrolaz, Gaël Panis, Bastien Casu, Silvia Ardissone, Laurence Degeorges, Martin Pilhofer, Patrick H. Viollier

## Abstract

The acquisition of multi-drug resistance (MDR) determinants jeopardizes treatment of bacterial infections with antibiotics. The tripartite efflux pump AcrAB-NodT confers adaptive MDR in the non-pathogenic α-proteobacterium *Caulobacter crescentus* via transcriptional induction by first-generation quinolone antibiotics. We discovered that overexpression of AcrAB-NodT by mutation or exogenous inducers confers resistance to cephalosporin and penicillin (β-lactam) antibiotics. Combining two-step mutagenesis-sequencing (Mut-Seq) and cephalosporin-resistant point mutants, we dissected how TipR uses a common operator of the divergent *tipR* and *acrAB-nodT* promoter for adaptive and/or potentiated AcrAB-NodT-directed efflux. Chemical screening identified compounds that interfere with DNA-binding by TipR or induce its dependent proteolytic turnover. We found that long-term induction of AcrAB-NodT disfigures the envelope and that homeostatic control by TipR includes co-induction of the DnaJ-like co-chaperone DjlA, to boost pump assembly and/or capacity in anticipation of envelope stress. Thus, the adaptive MDR regulatory circuitry reconciles drug efflux with co-chaperone function for trans-envelope assemblies and maintenance.

## INTRODUCTION

Bacterial multi-drug resistance (MDR) is jeopardizing treatment of bacterial infections with antibiotics (Karam et al., 2016). The MDR challenge is particularly grave for diderm bacteria that already possess an intrinsic MDR determinant: a protective outer membrane (OM) that prevents entry of soluble antibiotics. The OM of Gram-negative bacteria is typically structured as asymmetric lipid bilayer, with an inner leaflet of phospholipids and an outer leaflet containing lipo-polysaccharide (LPS), a charged glycolipid. By contrast, the inner (or cytoplasmic) membrane is a phospholipid bilayer, but both IM and OM flank the bacterial cell wall (CW, Figure 1A), an important shape-determining structure that is essential for life under most growth conditions. While the CW is the target of many natural antibiotics, the OM prevents these antibiotics from reaching the CW. Moreover, owing to its unusual lipid composition, the OM is also hydrophilic barrier for hydrophobic antibiotics(Nikaido and Pages, 2012; Ruiz et al., 2006).

**Figure 1:**
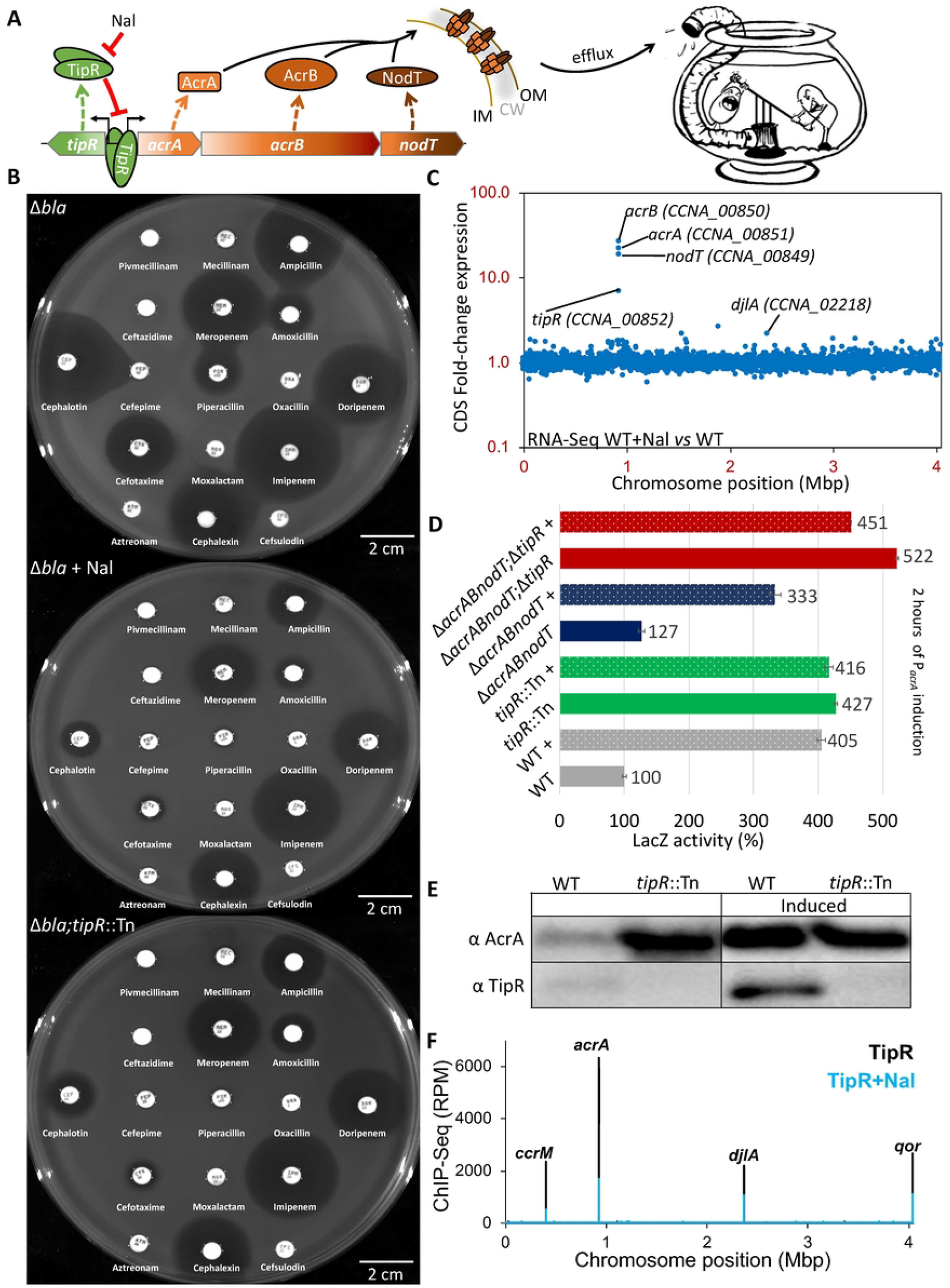
NAL-dependent de-repression of AcrAB-NodT and TipR confers adaptive β-lactam resistance. A) Schematic of the *tipR*/*acrAB-nodT* locus and the repression of the divergent promoter by TipR which is antagonized by the quinolone antibiotic nalidixic acid (Nal). The AcrAB-NodT components assemble into tripartite complex spanning the inner membrane (IM), cell wall (CW, grey shading) and outer membrane (OM) to expel diverse compounds as an efflux pump (as illustrated by the comic on the right). B) Antibiograms of *C. crescentus* strains on PYE. Antibiotic discs, from top left to bottom right: Pivmecillinam 20 µg, Mecillinam 10 µg, Ampicillin 100 µg, Ceftazidime 40 µg, Meropenem 10 µg, Amoxicillin 4 µg, Cephalothin 30 µg, Cefepime 30 µg, Piperacillin 100 µg, Oxacillin 5 µg, Doripenem 10 µg, Cefotaxime 30 µg, Moxalactam 30 µg, Imipenem 10 µg, Aztreonam 30 µg, Cephalexin 40 µg, Cefsulodin 30 µg. Nalidixic acid (Nal) induction performed at 20 µg/mL. C) Dot plot representation of the RNA-Seq analysis realised on *C. crescentus* NA1000 (WT). RNA was extracted from biological replicates of exponential phase cells harvested after induction with and without Nalidixic acid (Nal) 20 µg/mL for 30 minutes in PYE. Each dot represents the fold-enrichment of genes expression across the genome (represented as linear starting from the origin of replication) in the Nal induced condition. D) Activity of β-galactosidase of the P_*acrA*_-*lacZ* in different mutants of *C. crescentus*. The « + » indicates induction by Nalidixic acid 10 µg/mL for 2 hours. All levels are indicated in percentage of expression regarding the basal level of NA1000 (WT) without induction. E) Immunoblot anti-AcrA and anti-TipR performed on NA1000 (WT) and *tipR* mutant (*tipR*::Tn) cells with and without 2 hours of Nal 20 µg/mL induction. F) Representation of all TipR binding sites on the chromosome of *Caulobacter crescentus* (represented as linear starting from the origin of replication). The black line represents the Reads Per Million (RPM) obtained in the condition without induction while the blue line represent the RPM after 30 minutes of Nal 20 µg/mL treatment. Name above indicate the locus.

Additional protection against antibiotics that manage to enter the cell through OM pores comes from inducible envelope-spanning multi-drug efflux pumps, such as members of the tripartite resistance-nodulation-division (RND) transporter family (Lamut et al., 2019; Nikaido and Pages, 2012) that can expel a myriad of different noxious molecules (including antibiotics) from the cell using the proton motive force (PMF). Because the expression of many RND pumps is inducible by small molecules, they essentially function as adaptive MDR determinants. In the uninduced state, transcriptional repressors ensure low expression of efflux pumps by binding to their promoters to prevent firing. However, certain small molecule inducers can interact with the repressors, dislodging them from the promoter to relieve repression of the efflux pump gene (Ahmed et al., 1994; Li et al., 2015)(Figure 1A). Thus, these transcriptional repressors act as single component chemical sensors allowing a rapid protection by expulsion of toxic molecules that are encountered and produced, for example, by competing microbes.

The RND-type multidrug efflux pump AcrAB-TolC from the Gram-negative enterobacterium *Escherichia coli* has a wide substrate specificity, including clinically important antimicrobial compounds, dyes, fatty acids, monovalent and bivalent cationic lipophilic antiseptics and disinfectants, detergents and solvents (Li et al., 2015; Nikaido and Pages, 2012). The tripartite system is encoded in the *acrAB-tolC* operon that is repressed by the cis-encoded transcriptional repressor AcrR (Deng et al., 2013). We described a closely related RND efflux pump, AcrAB-NodT, that confers adaptive resistance because it is expressed from a promoter (P_*acrA*_) that is inducible by the first-generation quinolone nalidixic acid (NAL) and related molecules (Kirkpatrick et al., 2016; Kirkpatrick and Viollier, 2014). P_*acrA*_ is repressed by TipR, a TetR-like DNA-binding protein that is divergently encoded and expressed from a shared promoter region that spans 167 nucleotides between the predicted start codons of AcrA and TipR (Figure 1A).

TetR-type transcriptional repressors are characterised by an N-terminal DNA binding domain with a helix-turn-helix (HTH) motif and a C-terminal regulatory domain interacting with ligands (Ramos et al., 2005). Binding of an inducer to the C-terminal domain induces a conformational change that ultimately causes its dissociation from the target DNA, liberating the promoter to fire (Orth et al., 1998; Tiebel et al., 2000). TetR-like proteins are known to regulate a wide range of cellular functions, ranging from osmotic and general stress homeostasis, envelope maintenance, virulence gene expression, metabolism and synthesis of antimicrobial compounds, multi-drug resistance and expression of efflux pumps (Colclough et al., 2019; Cuthbertson and Nodwell, 2013; Ramos et al., 2005). Whether TipR also has such broad sensory and protective properties has not been explored.

Here we report that NAL-based induction of AcrAB-NodT in *C. crescentus* confers cross-protection against penicillins and cephalosporins, β-lactam antibiotics that target CW biosynthesis enzymes. We exploit this relationship in a deep mutational analysis that allowed dissecting the molecular determinants and mechanism responsible for TipR-dependent perception of various novel inducers identified in chemical screens. We also discovered that TipR not only acts in *cis* to control its own expression and that of AcrAB-NodT, but also targets a distant promoter that represses expression of DjlA, a DnaJ-like co-chaperone with a signature J-domain present in co-chaperones interacting with the DnaK protein folding chaperone (Castanie-Cornet et al., 2014; Mayer and Kityk, 2015). Our finding that envelope integrity is perturbed upon forced long-term over-expression of AcrAB-NodT and that DjlA augments AcrAB-NodT function, thus reconciles the genetically controlled co-induction of DjlA and AcrAB via TipR. This circuit design ensures a transient burst of efflux pump and co-chaperone co-expression to prevent efflux pump protein aggregation, while also augmenting efflux capacity.

## RESULTS

### NAL-induced adaptive resistance to β-lactams requires AcrAB-NodT

*C. crescentus* Δ*bla* mutant cells that lack the chromosomally-encoded metallo-β- lactamase (MBL) are unable to grow on plates supplemented with certain β-lactams, including the cephalosporin cephalothin (CEF, 10 µg/mL, CEF^10^) or the ureidopenicillin piperacillin (PIR, 40 µg/mL, PIR^40^). In an attempt to probe for determinants conferring β-lactam resistance in cells lacking MBL, we conducted a transposon (Tn) mutagenesis of Δ*bla* mutant cells, delivering the Tn by intergeneric conjugation from an *E. coli* donor that we sought to counter-selected on plates containing nalidixic acid (NAL, 20 µg/mL, NAL^20^). Unexpectedly, we noted that Δ*bla* cells grew on PIR^40^ or CEF^10^ plates supplemented with NAL^20^. NAL is not known to affect β-lactam resistance, but targets the A subunit of DNA gyrase (GyrA) in susceptible bacteria. However, *C. crescentus* is naturally resistant to NAL owing to the natural polymorphism F96D in GyrA (Kirkpatrick and Viollier, 2014). To test whether NAL affects β-lactam resistance in *C. crescentus*, we conducted Kirby-Bauer-based diffusion assays using β-lactam discs placed on plates with or without NAL^20^ that had been seeded with Δ*bla* cells. Indeed, we observed that NAL induces resistance of Δ*bla* cells to various cephalosporins (Figure 1B: CEF, cefotaxime and cephalexin) and penicillins (PIR, ampicillin, and amoxicillin), yet it provided little benefit to Δ*bla* cells against carbapenems (meropenem, imipenem and doripenem).

To dissect the genetic basis for the NAL-induced protection of these β-lactams, we mutagenized Δ*bla* cells with a *himar1* transposon (Tn) conferring kanamycin (KAN) resistance and counter-selected *E. coli* Tn-delivering donor cells with colistin or aztreonam (avoiding NAL) that C. crescentus is naturally resistant to. We selected for colonies on plates with KAN (20 µg/mL) and PIR^40^ and mapped the Tn insertions of two different mutant clones to the previously identified *tipR* gene (Figure 2A and Figure 2C). TipR is a cis-encoded repressor of the operon encoding the tripartite RND-type efflux pump AcrAB-NodT that is expressed from a NAL-inducible promoter (Kirkpatrick et al., 2016; Kirkpatrick and Viollier, 2014). We backcrossed the *tipR*::Tn mutation into Δ*bla* cells and compared the adaptive resistance conferred by NAL in Δ*bla tipR*::Tn double mutant cells to that of Δ*bla* parental cells by Kirby-Bauer antibiotic disc diffusion assays (Figure 1B) and, simply, by growth on plates containing CEF^10^ or PIR^40^ (Figure 2C). These assays revealed an identical β-lactam resistance spectrum between the Δ*bla* strain grown in the presence of NAL and the Δ*bla tipR*::Tn strain without NAL, indicating that NAL-responsiveness is lost in the absence of TipR (Figure 1B and Figure 2C).

**Figure 2:**
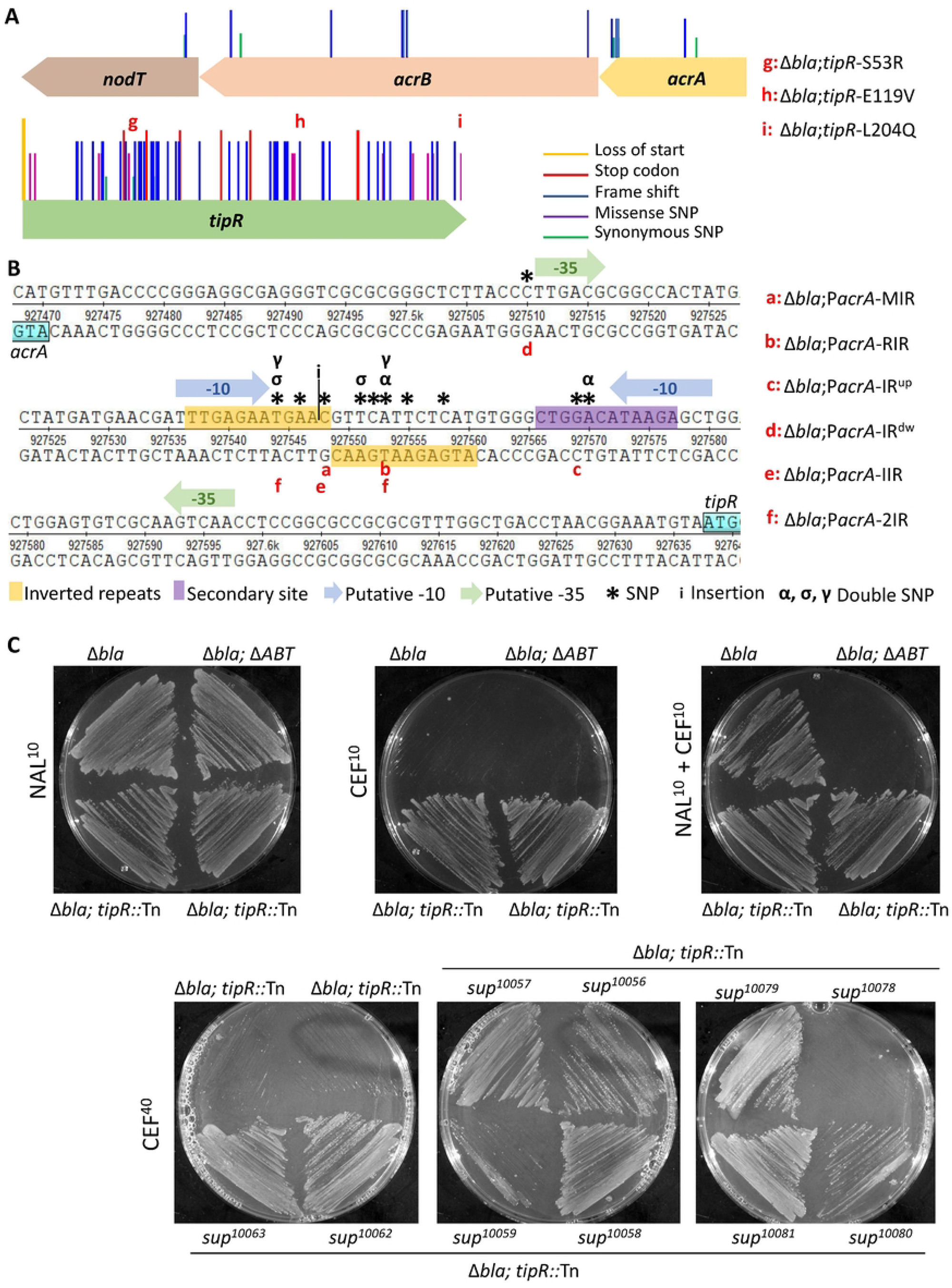
Deep mutational dissection of the *tipR*-*acrAB-nodT* locus. A) Positions of mutation in *tipR* and the *acrAB-nodT* operon conferring resistance to cephalothin 10 µg/mL to Δ*bla*. The list of mutations is a combination of sequences from the Mut-Seq experiment and single clone sequencing. The line represents, from the tallest to the smallest: the loss of start, new stop formation, missense mutations, frame shift and a synonymous variant. B) Intergenic region between *acrA* and *tipR*. The inverted repeats region is a hot spot for mutations that confer resistance to cephalothin 10 µg/mL to Δ*bla.* The * represent SNPs, (α, σ, γ) are double mutations per strain and (i) insertion. Position of the putative -10 and -35 sequences was predicted using SoftBerry (Solovyev, A Salamov (2011)). C) Resistance of Δ*bla* and various mutant derivatives to cephalothin or piperacillin. Strains were straked plates containing cephalothin 10 µg/mL (CEF^10^) or 40 µg/mL (CEF^40^), or piperacllin 40 µg/mL (PIR^40^) with or without nalidixic acid (10 µg/mL, NAL^10^) and grown for 72 hours at 30°C. Note that Δ*ABT* corresponds to the abbreviation for the Δ*acrAB-nodT* mutation.

RNA-Seq analysis of NAL-induced cells revealed a 20-fold induction of the *acrAB*-*nodT* transcript compared to the (uninduced) control state (Figure 2B, Figure S2, Table S1), as well as a 7-fold induction of the *tipR* transcript and a 2-fold induction of the transcript encoding the DnaJ-like co-chaperone DjlA (*CCNA_02218,* see below). Moreover, several transcripts encoding components of respiratory chain/cytochrome biosynthesis pathway (CCNA_01767, CCNA_2850, CCNA_3065) were also mildly upregulated. Thus, the transcriptional response to NAL predominantly affects the *acrAB-nodT* and *tipR* locus. No signature of a general transcriptional response to altered DNA supercoiling or DNA damage was apparent, a response that can occur in the presence of certain quinolone antibiotics. While the induction of the *djlA* transcript may point a direct interplay with TipR and AcrAB-NodT (see below), we reasoned that the weak upregulation of transcripts encoding respiratory proteins may reflect a response to increased consumption of the PMF from elevated AcrAB-NodT-mediated efflux.

To confirm that NAL relieves TipR-mediated repression of P_*acrA*_, we conducted β-galactosidase (LacZ)-based promoter probe assays (Figure 1D) in cells transformed with a P_*acrA*_-*lacZ* transcriptional reporter plasmid (pP_*acrA*_-*lacZ*) driving expression of a promoterless *lacZ* gene. In wild-type (*WT*) cells, the addition of NAL lead to a fourfold induction of P_*acrA*_-*lacZ* activity (405%) relative to the reference state without NAL (set to 100%). By contrast, P_*acrA*_-*lacZ* activity was 427% in *tipR*::Tn cells grown without NAL and remained unchanged upon addition of NAL. Matching results were obtained by monitoring the accumulation of AcrA by immunoblotting using antibodies to AcrA (Figure 1E and Figure S3A). AcrA abundance was strongly induced upon the addition of NAL in *WT* cells. By contrast, AcrA steady-state levels are elevated in *tipR*::Tn cells and not further augmented by the addition of NAL. We conclude that NAL-induction of P_*acrA*_ is likely governed through inactivation or removal of TipR.

Next, we monitored the induction of AcrA by NAL as a function of time (Figure S3B) and observed AcrA abundance to peak 20 minutes after the addition of NAL. Moreover, and consistent with the induction of *tipR* mRNA by RNA-Seq, immunoblotting using polyclonal antibodies to TipR revealed a congruent accumulation of TipR with AcrA during the time course with NAL (Figure S3B), suggesting that P_*acrA*_ and the *tipR* promoter (P*_tipR_*) are simultaneously de-repressed upon the addition of NAL. ChIP-Seq experiments conducted with polyclonal antibodies to TipR revealed that TipR binds to four chromosomal sites *in vivo*. Importantly, TipR occupancy of these sites is drastically reduced after exposing cells to NAL (for 30 minutes Figure 1F, Figure S2), indicating that NAL antagonizes TipR binding *in vivo*. These experiments further support the view that TipR represses P_*acrA*_ and P*_tipR_*, and that NAL treatment directly or indirectly acts on TipR to induce these two promoters and force their release from TipR repression. The coordinated induction of AcrAB-NodT and the TipR proteins by NAL points towards a homeostatic control mechanism in which the adaptive resistance to β-lactam antibiotics is conferred by induction of AcrAB-NodT, followed by TipR synthesis to re-establish the repressed ground state once the inducers have been expelled by AcrAB-NodT.

As the NAL-induced adaptive β-lactam resistance is easiest to score in Δ*bla* cells that are hypersensitive to these antibiotics, we confirmed that the adaptive resistance is lost in Δ*bla* Δ*acrAB-nodT* cells (Figure 2C and Figure S1A). While this result indicates that the resistance is mediated by induction of AcrAB-NodT, we found that the low-level expression of AcrAB-NodT in the uninduced state can confer resistance to the β-lactam cefepime, an effect that can also be seen to a lesser extent for cephalothin and cefotaxime (Figure S1B). With regards to cephalothin, we conducted a second step selection with Δ*bla tipR*::Tn cells to isolate mutants with a high-level of cephalothin resistance (CEF^40^), exceeding the CEF^10^ resistance of Δ*bla tipR*::Tn cells. In fact, we isolated Δ*bla tipR*::Tn gain-of-function mutants on CEF^40^ plates (cephalothin 40 µg/mL) harbouring single missense mutations in *acrB* (Q177P, L241P, Q528H, P565A, E566K, N676Y, I762F; Figure 2C, Figure S2B), presumably due to increased specificity, capacity or abundance to expel cephalothin (Figure 2C). These experiments further support the role of AcrAB-NodT in adaptive resistance to cephalothin, and further point to AcrB as a key plasticity determinant enabling high-level MDR.

### Mut-Seq and genetic dissection of TipR and its target

Having established the chemical-genetic relationship between AcrAB-NodT and cephalothin, we exploited the CEF^10^ selection to disentangle how TipR’s interaction with its target is modulated. To this end, we first plated Δ*bla* cells on agar containing CEF^10^ to isolate spontaneous point mutations conferring loss-of-function in *tipR*, or ptentially its (unknown) target sequence in P_*acrA*_. We then collected several hundred CEF^10^-resistant Δ*bla* mutant cells per plate, pooled them, PCR amplified *tipR* and *acrAB*-*nodT* from the mutant pool and deep-sequenced the PCR products. Sequence analysis (Figure 2A, Table S2) of this Mut-Seq dataset indeed revealed missense, non-sense and frameshift mutations scattered throughout *tipR*, with slight hotspot in the region predicted to encode the N-terminal DNA-binding domain. The fact that nonsense and frameshift mutations are also found in the C-terminal half of the protein indicates that these residues are also important for TipR function. Interestingly, we also found several (gain-of-function) mutations within the *acrAB*-*nodT* coding sequence that presumably increasing the stability, structure and/or affinity of AcrAB-NodT components towards CEF (matching the CEF^40^ experiments described above).

Next, we isolated individual CEF^10^-resistant Δ*bla* mutant clones from this pool and sequenced their genomes. Three TipR missense mutants were chosen for further study, each encoding a different mutation in TipR: S53R (in the DNA-binding domain), as well as E119V and L204Q (Figure 2A, corresponding to mutants Δ*bla tipR*-*S53R,* Δ*bla tipR*-*E119V* and Δ*bla tipR*-*S53R*) in the central region and C-terminal domain. We confirmed the strong elevation of P_*acrA*_ activity in these mutants by P_*acrA*_-*lacZ*-based promoter probe assays (Figure S4).

While these missense mutants in TipR are thus indeed loss-of-function mutations, we also to obtain clones with mutations in the operator site for TipR (or in the promoter of *tipR*, P*_tipR_*, since *tipR* and *acrAB-nodT* are transcribed divergently from the same intergenic region, Figure 2B) that would prevent TipR from binding and repressing P_*acrA*_. Since the binding site of TipR is smaller than the 615-nucleotide *tipR* gene, these loss-of-function mutations in P_*acrA*_ would likely surface less frequently than nonsense mutations in *tipR*. Nonetheless, our genome sequencing of CEF^10^-resistant mutants revealed two strains with mutations in P_*acrA*_, (strains Δ*bla* P_*acrA*_-*RIR* and Δ*bla* P_*acrA*_-*IR^up^*), attesting to the strength of the selection. To favour the enrichment of loss-of-function mutations in P_*acrA*_ (Figure 2B) that enable a throurough genetic dissecting the TipR target sequence, we modified our starting genotype for a revised screen. In fact, we conducted the CEF^10^-resistant selection with Δ*bla* cells carrying a multi-copy plasmid expressing TipR (pSRK-Gm-*tipR*) to eliminate mutants with a dysfunctional *tipR* gene that would otherwise surface. With this strategy, we uncovered strains with additional P_*acrA*_ mutations, 80% of these mutations reside in a region of dyad symmetry comprising an eleven base pair inverted repeated (IR, 5’-TGAGAATGAAC-3’, Figure 2B), two adjacent mutations are eleven nucleotides upstream of the IR (IR*^up^* mutation) and another mutation lies thirty-three nucleotides downstream of the IR (mutation IR*^dw^*).

Lastly, we chose the following IR mutant strains for follow-up confirmation studies: Δ*bla* P_*acrA*_-MIR (Middle Inverted Repeat), Δ*bla* P_*acrA*_-RIR (Right Inverted Repeat), Δ*bla* P_*acrA*_-2IR (2 mutations Inverted Repeat) and Δ*bla* P_*acrA*_-IIR (Insertion Inverted Repeat, Figure 2B), as well as one with a mutation upstream of the IR (Δ*bla* P_*acrA*_-IR*^up^*) and one downstream of it (Δ*bla* P_*acrA*_-IR*^dw^*, Figure 2B). Immunoblotting using antibodies to AcrA and TipR confirmed that AcrA is overexpressed in all selected mutants compared to the Δ*bla* parent (Figure S5A-D), explaining why these mutants grew on CEF^10^ plates. TipR was also overproduced in all mutants, except for strains Δ*bla* P_*acrA*_-IR*^up^* and Δ*bla tipR*-*L204Q* (Figure S5A-B) in which the level was similar to the basal amount of the Δ*bla* parental strain. The addition of NAL led to the induction of TipR in these mutants, albeit to lower steady-state levels compared to Δ*bla* cells, whereas AcrA levels were not substantially affected. This result suggests that the effect of the mutation differs for P*_acrA_ versus* P*_tipR_*, and/or that NAL affects the abundance of TipR through a post-translational mechanism. Below we provide additional evidence for the latter model.

In summary, the strong selection for CEF^10^-resistance in Δ*bla* cells unveiled mutational hotspots in the region predicted to be required for DNA-binding by TipR, but also in the IR of in the suspected promoter target of TipR.

### Defining the TipR target sequence and residues required for DNA-binding

We used electrophoretic mobility assays (EMSAs) to confirm that TipR binds the IR *in vitro*. In these experiments, a Cy5-labelled P_*acrA*_ probe centred on the IR was mixed with recombinant (untagged) TipR purified in two steps from an *E. coli* overexpression strain (see Materials and Methods). We observed a retardation of the labelled probe by TipR in a concentration-dependent fashion, even in the presence of non-specific competitor DNA (calf thymus DNA) provided in excess (Figure 3A). EMSA experiments with *E. coli* extracts containing (*WT*) TipR also revealed retardation of Cy5-labelled P_*acrA*_ by compared to *E. coli* extracts without TipR. Next, we probed *E. coli* extracts each containing a different TipR mutant described above (E119V, S53R or L204Q) for binding of P_*acrA*_ by EMSA. While the L204Q variant retards P_*acrA*_ akin to *WT* TipR, the E119V or S53R mutations abolish P_*acrA*_ binding (Figure 3A). Immunoblots conducted with *C. crescentus* cell extracts revealed the E119V and S53R variants to accumulate to the same steady-state level as *WT* TipR upon NAL induction (Figure S5), indicating that the E119V and S53R mutations promote DNA binding. By contrast, the abundance of TipR-L204Q is severely reduced, even in the presence of NAL (Figure S5A), suggesting that this mutation interferes with protein stability rather than with DNA-binding activity. Indeed, antibiotic chase experiments revealed a reduced half-life of TipR-L204Q (Figure 3B).

**Figure 3:**
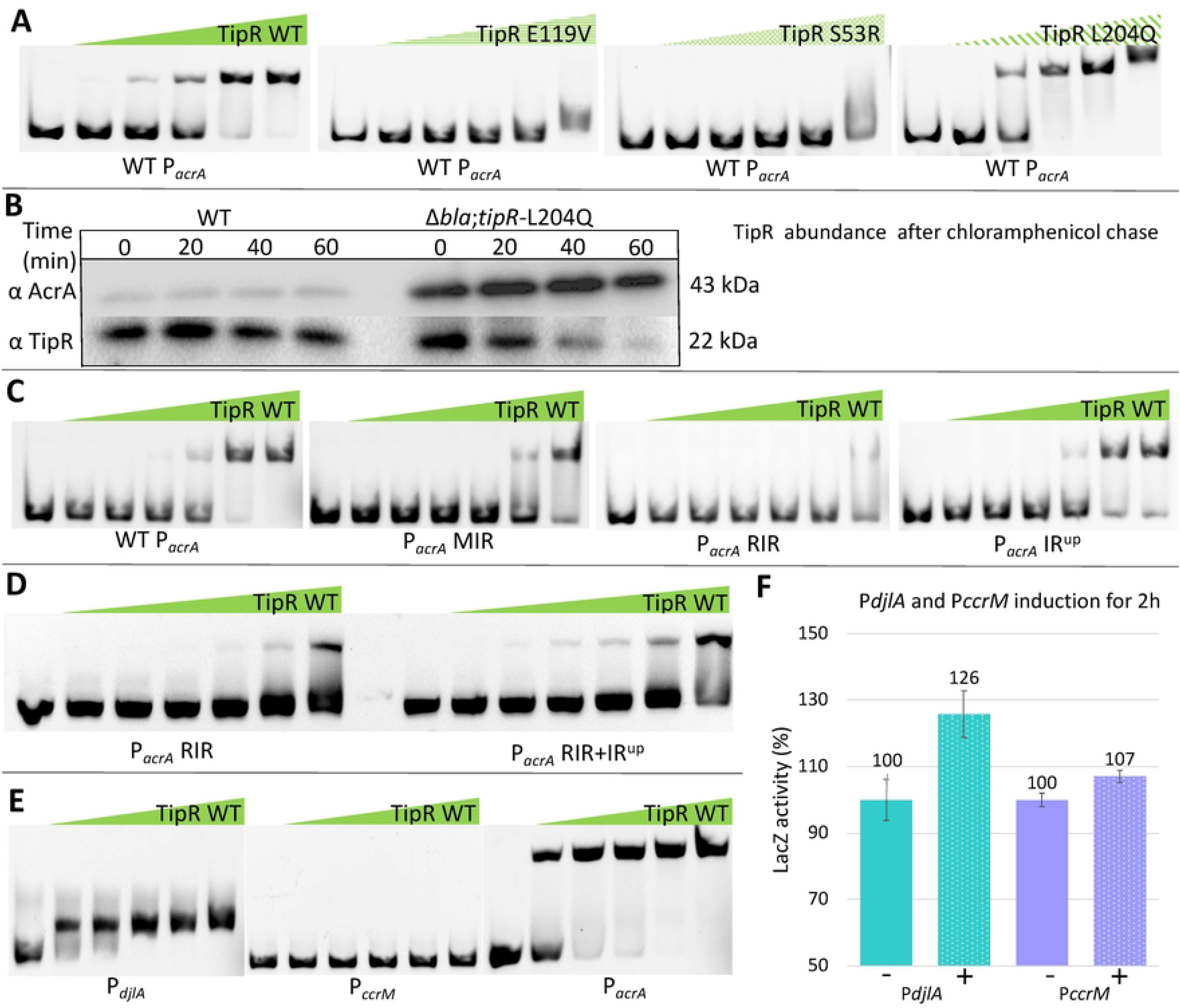
Determinants in TipR and P_*acrA*_ required for repression. A) EMSA with 200 ng of a Cy5-labelled *acrA* promoter (P_*acrA*_) probe in the presences of increasing concentration of cell extract from *E. coli* overexpressing different mutated TipR protein. Concentration of protein start at 0,031 µM to 16 µM. B) Western blot showing the result of a 1h chase experiment using chloramphenicol 5 µg/mL on exponential cells of NA1000 (WT) and Δ*bla*;*tipR*-L204Q in PYE. C) EMSA with 200 ng of different Cy5-labelled P_*acrA*_ mutants analysed for retardation by gradual concentration of purified TipR protein. Concentration of protein start at 0,0156 µM to 16 µM. D) EMSA with 100 ng of Cy5-labelled *acrA* with gradual concentration of heparin-purified TipR protein. Concentration of protein start at 0,25 µM to 8µ M. E) EMSA with 100 ng of Cy5-labelled *djlA, ccrM* and *acrA* promoters (P*_djlA_*, P*_ccrM_* and PacrA) with gradual concentration of purified TipR protein at concentrations of 1.125 µM to 18 µM. F) Activity of β-galactosidase of the P*_djlA_*-*lacZ* or P*_ccrM_*-*lacZ* in *WT C. crescentus*. The « + » indicates induction by Nalidixic acid 10 µg/mL for 2 hours. All levels are indicated in percentage of expression regarding the basal level of NA1000 (WT) without induction.

Having identfied determinants in TipR that promote DNA-binding, we next determined the effects of the isolated promoter mutations. To this end, P_*acrA*_ promoter mutants with alterations in the IR were labelled with Cy3 and incubated with purified WT TipR in EMSAs (Figure 3C). While P_*acrA*_-RIR is no longer retarded by TipR, we only observed a partial loss of binding by TipR to P_*acrA*_-MIR. As expected TipR still retarded the P_*acrA*_-IR^up^ probe that carries a mutation outside the IR. Thus, these EMSAs validate the requirement of the IR as the likely TipR docking site in P_*acrA*_. With these results, we investigated further how P_*acrA*_-IR^up^ can cause overexpression of the pump without crippling DNA-binding of TipR. EMSAs with a new Cy5-P_*acrA*_ probe combining the P_*acrA*_-RIR mutant (that impairs binding of TipR) with the P_*acrA*_-IR^up^ mutation, EMSA (Figure 3D) revealed that P_*acrA*_-IR^up^ increased the affinity of TipR for the P_*acrA*_-RIR probe suggesting that P_*acrA*_-IR^up^ mutation creates a secondary docking site upstream of the IR.

Indeed, the consensus target sequence derived for TipR (Figure S7) resembles the motif created by the P_*acrA*_-IR^up^ mutation. In search for additional binding sites of TipR, we conducted ChIP-Seq experiments and found that *in vivo* TipR binds four sites on the chromosome: the promoter of *acrAB-nodT* and *tipR*, the promoter of the *djlA* gene predicted to encode a DnaJ-like co-chaperone, as well as a site upstream of the DNA methyltransferase gene *ccrM* and another in between the *qor* and *rho* genes that face each other. The RNA-Seq experiments described above confirmed the NAL induction of three genes neighbouring two of the TipR binding site (*acrAB-nodT*, *tipR* and *djlA*) and ChIP-Seq experiments conducted with NAL-treated cells revealed that the binding of TipR to all four chromosomal sites is impaired *in vivo* (Figure 1F, Figure S6). However, the binding of TipR to P_*acrA*_ was more efficient than to the other sites, suggesting that the IR may represent the preferred recognition sequence of TipR. Nonetheless, we scanned the other target sites for sequences comparable to the IR and detected one upstream of *djlA* and of *ccrM,* delivering the consensus sequence 5’-WTGaGWMtGAWC-3’ by MEME analysis (Figure S7). Using this consensus, we analysed the intergenic sequence between the *qor* and *rho* genes and found a single sequence that could explain the presence of TipR at this location, however the configuration of those genes facing each other make a TipR-dependant regulation unlikely. We then tested the binding ability of TipR to *djlA* and *ccrM* promoters *in vitro* by EMSA using purified recombinant TipR. As seen in Figure 3E, TipR was able to retard the *djlA* promoter, but not the promoter of *ccrM*. Moreover, LacZ-based promoter probe assays with P*_djlA_*-*lacZ* and P*_ccrM_* -*lacZ* reporters revealed an induction of P*_djlA_* in *WT* cells in response to NAL, but no significant induction of P*_ccrM_*-*lacZ* (Figure 3F). Since TipR binds P*_ccrM_* weakly *in vivo*, likely because of the divergence of its IR from the preferred TipR target sequence, the weak interaction might result in a TipR-promoter complex too fragile to be maintained during electrophoresis. Moreover, in the absence of a significant induction of transcription by NAL in our RNA-Seq experiment (Figure 1C), we speculate that TipR is a poor repressor of P*_ccrM_ in vivo*, yet it clearly down-regulates P*_djlA_*.

### Screening for chemical inducers of P_*acrA*_

Having identified the IR of P_*acrA*_ as the target of TipR’s DNA-binding activity, we engineered a reporter for rapid screening of chemical libraries for compounds that induce P_*acrA*_ (independent of requirement of AcrAB-NodT efflux function as in CEF^10^ resistance screen above). To this end, we fused to the promoterless *nptII* kanamycin resistance gene to P_*acrA*_, creating a convenient readout for molecules like NAL that can activate P*_acrA_ in vivo* by probing for kanamycin resistance. We transformed the resulting P_*acrA*_-*nptII* promoter probe reporter plasmid (pP_*acrA*_-*nptII*) into *WT* cells, embedded the resulting reporter cells in soft agar containing kanamycin (10 µg/mL, KAN^10^) overlaid on KAN^10^ plates and then spotted 4µL drops of each compound (at 10 µM) of the Maybridge chemical library for inducers of P_*acrA*_ on these seeded indicator cells. Small molecule inducers were identified by their ability to promote growth of cells around the disc owing to activation of P_*acrA*_-*nptII* by the compound(s) (Figure S8). This chemical screen along with tests of candidate compounds reported in the literature for other systems, unearthed 14 different inducers P_*acrA*_-*nptII* (Figure 4A and Figure S8) that were subsequently grouped into two inducer classes based upon pP_*acrA*_-*lacZ* inducer strength as determined by LacZ activity measurements. One group of strong inducers (triggering P_*acrA*_-*lacZ* activity exceeding 230% relative to *WT*) includes the quinolones NAL, flumequine (FLU) and sparfloxacin; quinolone precursors such as chloroquine and dyes such as rhodamine 6G (Rh6) and Crystal Violet (CV). The weak inducers include the dye Malachite Green (MG), DNA intercalants such as Acridine Orange (AO), Ethidium Bromide (EtBr) and SYBR safe, but also various other compounds such as glafenine and doxylamine. We also noticed that dopamine and aminolevulinic acid fail to induce P_*acrA*_-*lacZ* after 3h, however they allowed growth on kanamycin plates with P_*acrA*_-*nptII* after overnight incubation (Figure 4A and Figure S8), suggesting that they act slowly or that they counteract kanamycin in another way.

**Figure 4:**
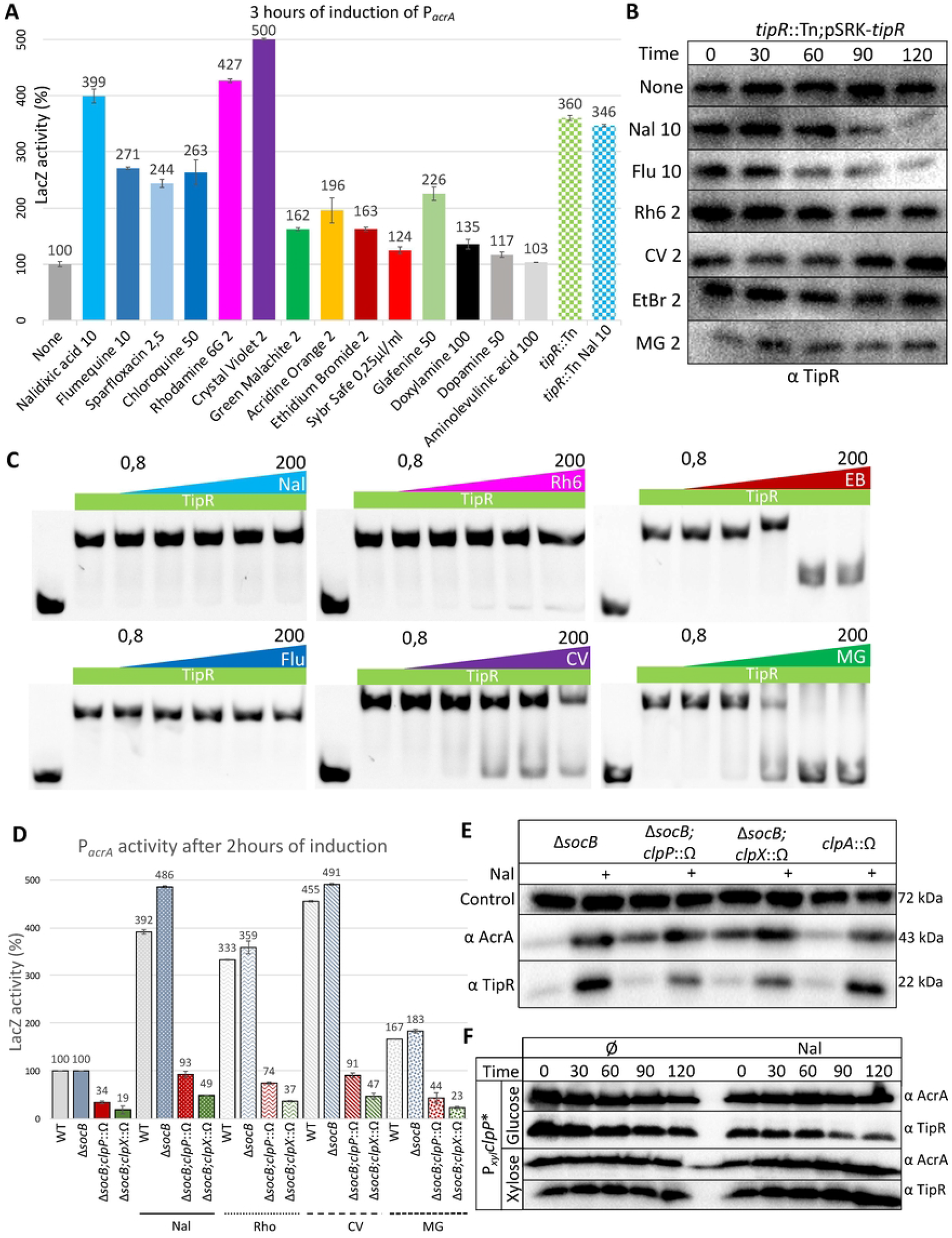
Different chemicals relieve repression of P_*acrA*_ by TipR. A) Expression of β-galactosidase assay using the P_*acrA*_-*lacZ* in NA1000 after 3 hours of induction by different compounds. All concentrations are in µg/mL. All levels are indicated in percentage of expression regarding the basal level of the control: NA1000 (*WT*) without induction. B) Immunoblot showing the TipR abundance during a 2 hour chase experiment with chloramphenicol 5 µg/mL on exponential *tipR*::Tn;pSRK-*tipR* cells grown in PYE with and without selected inducers: Nalidixic acid (Nal 10 µg/mL), Flumequine (Flu, 10 µg/mL), Rhodamine 6G (Rh6, 2 µg/mL), Crystal violet (CV, 2 µg/mL), Ethidium bromide (EtBr 2, µg/mL) and Malachite Green (MG, 2 µg/mL). C) EMSA using 200 ng of Cy5 tagged DNA of the promoting region of *acrA* with 8 µM of heparin purified TipR, supplemented with an increasing concentration of selected inducers. All quantities expressed in µg/mL. D) β-galactosidase measurements in *C. crescentus WT* and different mutants containing P_*acrA*_-*lacZ* before or after induction with nalidixic acid (Nal, 10 µg/mL), Rhodamine 6G (Rho,2 µg/mL Rho), Crystal Violet (CV 2 µg/mL) and Malachite Green (MG, 2 µg/mL) for 2 hours. All measurements are expressed as percentage of expression relative to basal activity of the control: *WT* or Δ*socB* cells before induction. E) Immunoblots on cell extracts from different protease mutants performed with anti-TipR and anti-AcrA antibodies. All inductions (+) were performed after 2 hours of induction with 10 µg/mL of Nalidixic acid (Nal). Shown as loading control are blots with antibodies to CCNA_00163. F) Immunoblot blot following a 2-hour chase experiment with chloramphenicol (5 µg/mL) on exponential cells of *WT* with a *clpP*\* inserted at the P*_xyl_* locus. Cells were grown in PYE, followed by the addition of glucose or xylose, and subsequently the inducer was added: Nalidixic acid (Nal 10 µg/mL) and finally protein synthesis was blocked with chloramphenicol.

Some of the identified molecules are known substrates of the enterobacterial AcrAB-TolC RND pump and would thus likely be exported to reduce the active concentration in cells. Hence, we wondered if induction of P_*acrA*_ would be accentuated in cells lacking AcrAB-NodT (Figure S9A) that can no longer expel these molecules, resulting in their accumulation to higher intracellular levels. While the level of P_*acrA*_ induction with compounds like NAL or EtBr was similar in *WT versus* Δ*acrAB-nodT* cells, AO clearly showed a 4-fold elevated level of induction compared to the 2-fold induction in *WT* cells. Thus, the intracellular concentration modulates the level of *acrAB-nodT* expression. To eliminate the possibility that the compounds cause a non-specific global increase of gene expression or LacZ enzymatic activity, we tested P*_bla_*-*lacZ* reporter expressing LacZ expressed from the constitutive promoter of the *C. crescentus* metallo-β-lactamase gene *CCNA_02223* and found that these compounds did not alter LacZ activity from P*_bla_*-*lacZ* (Figure S9B).

### Different inducers control TipR at the post-translational level

To determine if the new inducers act in the same way on TipR as NAL, we first explored if they affect TipR stability *in vivo*. To this end, we uncoupled TipR synthesis from its inducible transcription by expressing it from the IPTG-controllable P*lac* promoter (of *E. coli*) on plasmid pSRK-*tipR* in *tipR*::Tn cells. AcrAB-NodT is still inducible from P_*acrA*_ in these cells, but TipR synthesis is independent from the tester compounds such as NAL, Rh6 and MG (Figure S10A-B). We then compared the half-life of TipR in these cells after exposing them to the chemical inducers. To stop translation of TipR, we treated cells with high levels of the protein synthesis inhibitor chloramphenicol, a phenicol antibiotic that is not a substrate of the AcrAB-NodT efflux pump. In the presence of the quinolones NAL or FLU, TipR is rapidly turned over (Figure 4B), reminiscent of the instability observed for TipR-L204Q. Such an induced instability was not observed with the other P_*acrA*_-activating compounds.

Next, we examined the effect of these compounds on the DNA binding ability of purified TipR by EMSA (Figure 4C). While at physiologically relevant concentrations, NAL or FLU did not perturb retardation of the Cy-labelled P_*acrA*_, other inducers such as EtBr or MG interfered with binding of TipR to P_*acrA*_. Rh6 and CV affect TipR binding in the same manner, but less efficiently *in vitro* compared to EtBr or MG. To assess the specificity of these compounds on TipR, rather than interfering with protein-DNA interactions in a general fashion, we conducted control EMSA using the histone-like IHF protein and its target DNA sequence *attR* (Figure S11) as probe. Increasing concentration of CV did not prevent IHF binding even at a concentration sufficient to change the charge of the probe resulting in its upward migration in the gel during electrophoresis. Therefore, CV, Rh6, EtBr and MG apparently interfere with TipR’s ability to bind the IR in P_*acrA*_, explaining the de-repression of P_*acrA*_ by these compounds *in vivo*.

To rule out the possibility that induction of AcrAB-NodT by NAL or FLU acts through the SOS (DNA damage) response by corrupting DNA gyrase (encoded by *gyrAB*) as in *E. coli*, we confirmed that inhibition of GyrA by the fluoroquinolone ciprofloxacin (CIP) or of GyrB by the aminocoumarin novobiocin (NOV) alone did not induce of P_*acrA*_ (Figure S12). Moreover, NAL and CIP did not synergise to enhance P_*acrA*_ induction, yet NAL and NOV together resulted in elevated P_*acrA*_-*lacZ* induction. Consistent with these findings, we next assayed P_*acrA*_-*lacZ* activity in *WT* cells expressing an additional GyrA variant: GyrA from *Brucella melitensis* or a mutant variant of *C. crescentus* GyrA, GyrA*(F96D) that is NAL-sensitive and causes DNA damage and the SOS response in the presence of NAL. *C. crescentus WT* cells expressing either of these GyrA variants from pMT335 still induce P_*acrA*_ upon the addition of NAL (Figure S12), even more efficiently than *WT* cells. In fact, the level of NAL-induction in cells expressing the NAL-sensitive forms of GyrA attains the level of induction when NAL and NOV are added jointly to *WT* cells, indicating that Gyrase inhibition can enhance AcrAB-NodT induction, but only after repression of P_*acrA*_ by TipR has been relieved by the addition of NAL.

Prompted by the observation that, at physiological concentrations, NAL and FLU reduce the half-life of TipR without interfering with TipR binding to P*_acrA_ in vitro* in EMSAs (Figure 4B), we speculated that a transient destabilization by a dedicated protease could account for the induction of AcrAB-NodT. Using a candidate approach, we tested for protease(s) conferring instability of TipR by immunoblotting using antibodies to TipR and AcrA (Figure S13). While no significant changes in AcrA or TipR steady-state levels was observed in *ftsH* and *lon* mutant cells compared to *WT* cells, loss of ClpP or ClpX led to an accumulation of AcrA and a reduction in TipR levels in the absence of NAL. Indeed, previous work showed that AcrA and AcrB are substrates of the ClpP degradation machinery (Bhat et al., 2013). Yet, AcrA and TipR are still inducible upon exposure of *clpP* and *clpX* mutant cells to NAL (Figure 4E and Figure S13), but promoter probe assays using the P_*acrA*_-*lacZ* reporter revealed a strong reduction of LacZ-induction when *clpP* and *clpX* mutant are exposed to NAL, Rh6, CV or MG compared to *WT* cells (Figure 4D). Taken together our findings suggests that the ClpXP protease promotes high level of induction of *acrAB-nodT* and *tipR*, whereas inactivation of the *clpA* chaperone gene in *WT* cells did not cause a major AcrA or TipR abundance before after induction with NAL (Figure 4E).

To test if ClpP controls the half-life of AcrA and TipR, we conducted antibiotic chase experiments in cells expressing a dominant negative version of ClpP (ClpP*) in which the catalytic serine is mutated to alanine (S11A, Figure 4F). This ClpP* variant is expressed from the xylose-inducible P*_xyl_* promoter at the *xylX* locus (*xylX*::P*_xyl_*-*clpP*\*). Immunoblotting using antibodies to TipR and AcrA revealed that TipR is turned over normally upon the addition NAL in the presence of glucose (to repress expression of ClpP* from P*_xyl_*). However, when cells are grown in PYE containing xylose to induce of the dominant negative ClpP* variant), TipR stability is increased. Lastly, to demonstrate the interaction of TipR and AcrA with ClpXP biochemically, we used a GFP Trap matrix to pull down a ClpX-YFP fusion protein from lysates prepared from *WT xylX:.Pxyl-ClpX-YFP* cells before or after induction with NAL. Immunoblotting revealed the presence of TipR and AcrA in the pulled down material (Figure S14) regardless of the presence of NAL.

In summary, while some compounds dislodge TipR from the IR sequence, others other like first generation quinolones act to destabilize TipR. The ClpXP protease confers the instability of TipR in the presence of NAL and it also reduces TipR and AcrA steady-state levels in the absence of NAL. Lastly, we discovered that ClpXP is required for efficient transcriptional induction of *acrAB-nodT*.

### Envelope defects caused by AcrAB-NodT overexpression

Because *tipR*::Tn or NAL-treated *WT* cells grow slower than untreated *WT* cells, we imaged them by phase contrast microscopy and discovered that the frequently had envelope blebs and other envelope deformations. These blebs were seen less frequently when AcrAB-NodT induction occurred through NAL, than through inactivation of TipR or through simple AcrAB-NodT overexpression form a plasmid (Figure 5A, red arrows). To determine if this effect requires the efflux activity of AcrAB-NodT, we exposed these cells to the efflux pump inhibitor 1-(1-Naphthylmethyl)-piperazine (NMP) and still observed these envelope deformations (Figure 5A), suggesting that the assembly of this trans-envelope structure can disfigure cells. To investigate the nature of those perturbations in detail, we imaged cells by cryo-electron tomography (cryoET). We imaged Δ*bla tipR*::Tn cells and Δ*bla* cells carrying a plasmid containing *acrAB-nodT* under the control of an IPTG inducible promoter (pSRK-*acrAB-nodT*) and, in order obtain the maximal (conditional) overexpression of AcrAB-NodT, we also induced Δ*bla* cells harbouring pSRK-*acrAB-nodT* with IPTG and NAL^10^ (to induce AcrAB-NodT from the chromosome). The electron cryo-tomograms of these cells (Figure 5B) showed an intact inner membrane and peptidoglycan layer. However, the OM and the S-layer are highly aberrant, irregularly spaced (i.e. detached from the inner membrane) and crooked with major deformations including membrane blebs and invaginations, sometimes harbouring evaginated OM vesicles (Figure 5C, Figure S15). While the precise content and origin of those vesicles remains unknown, we suspect that they contain phospholipid-and/or LPS-derived OM material and/or soluble periplasmic content. Consistent with the observed envelope irregularities, we found that *tipR*::Tn cells are more susceptible peptidoglycan-targeting antibiotics than *WT* cells, for example vancomycin, teicoplanin, bacitracin and fosfomycin (Figure S16). Those molecules are known to be poorly efficient against Gram-negative bacteria due to the OM which prevent their entry into the cells (Muheim et al., 2017), suggesting that the observed deformations and blebs lead to higher cell permeability. By contrast, *tipR*::Tn cells are more resistant to the quinolones sparfloxacin and ciprofloxacin as well as to the macrolide antibiotics erythromycin and azithromycin, likely owing to increased efflux conferred by AcrAB-NodT overexpression.

**Figure 5:**
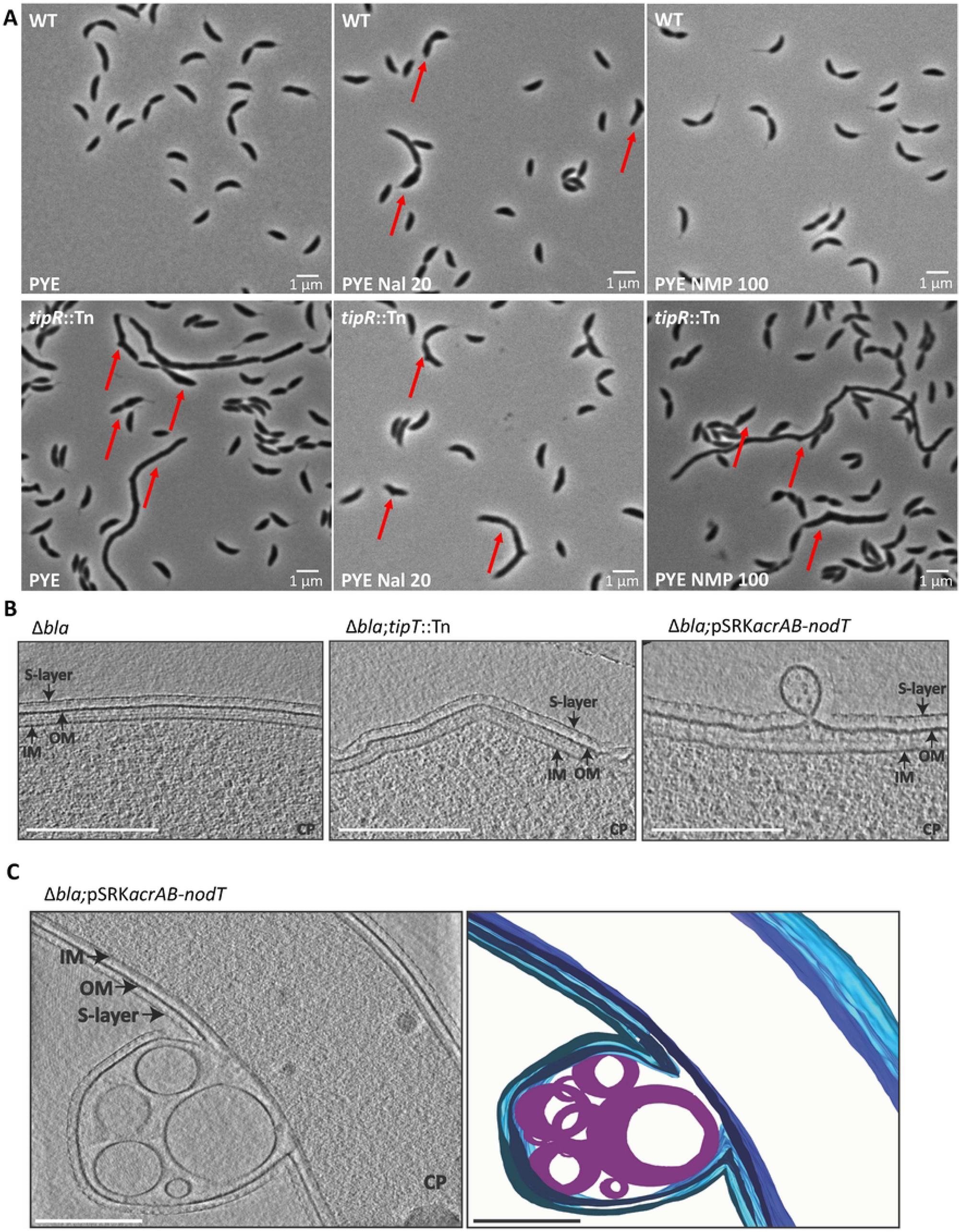
Envelope defects caused by long-term overexpression of AcrAB-NodT. A) Light microscopy (phase contrast) images of NA1000 (*WT*) and *tipR*::Tn cells during exponential growth in PYE in the presence or absence of Nalidixic acid (Nal, 10 1-(1-Naphtylmethyl)-piperazine (NMP quantities are indicated. B) Cryo-ET images *C. crescentus* Δ*bla* cells (left), Δ*bla*; *tipR*::Tn (center) and Δ*bla*; pSRK-*acrAB*-*nodT* (right). Scale bar, 250 nm. IM, inner membrane; OM, outer membrane; CP, cytoplasm. C) ECT images (left), 3D rendered image (center) and a superposition of both images (right) of *C. crescentus* Δ*bla*;*tipR*::Tn cells (A) and Δ*bla*;pSRK-*acrAB*-*nodT* (B), imaged by conventional cryo-ET. Scale bar, 250 nm. IM (dark blue), inner membrane; OM (blue), outer membrane; S-Layer (cyan); CP, cytoplasm; Lipid droplet (purple).

### The DjlA co-chaperone enhances AcrAB-NodT

Knowing that massive overexpression of AcrAB-NodT alone can strain envelope integrity and structure, while short-term induction of AcrAB-NodT and DjlA with NAL is less severe, we interrogated the possible role in DjlA co-expression with AcrAB-NodT. Specifically, we asked whether the DjlA co-chaperone augments AcrAB-NodT efflux activity, assembly and/or mitigates massive envelope damage. We first examined whether DjlA overexpression or inactivation affects the induction of P_*acrA*_ or TipR and AcrA by NAL. Neither P_*acrA*_-*lacZ*-activity, nor AcrA or TipR abundance was affected as a function of DjlA, regardless of whether NAL was present or not (Figure 6A, 6D and Figure S17). However, when we probed for AcrAB-NodT-mediated efflux activity, we found that ectopic expression of DjlA enhances resistance to several β-lactam antibiotics belonging to the cephalosporin and penicillin class that we showed above are substrates of AcrAB-NodT (Figure S18). To assess if DjlA promotes AcrAB-NodT -dependent efflux to other types of efflux substrates, we conducted survival (efficiency-of-plating, EOP) assays on plates containing EtBr^6^ (EtBr, 6 µg/mL) and found that cells ectopically expressing DjlA (from pMT335-*djlA*) exhibited an elevated EOP compared to *WT* cells with the empty vector (Figure 6B). The boosting effect of pMT335-*djlA* on survival on EtBr^6^ plates was also observed for Δ*djlA* cells, but no longer for Δ*acrAB-nodT* cells, indicating that the activity or assembly of the pump, and thus the ability to expel EtBr, is enhanced by DjlA. In support of this view, Δ*djlA* cells exhibit a reduction in EOP compared to *WT* cells on plates containing EtBr^4^ (Figure 6C). Interestingly, forcing overexpression of AcrAB-NodT in Δ*djlA* cells by introducing the *tipR*::Tn mutation or by the addition of NAL, can mitigate the reduced EOP on EtBr^4^ (EtBr, 4 µg/mL) plates, likely because AcrAB-NodT levels are no longer limiting for expulsion of EtBr to enable growth.

**Figure 6:**
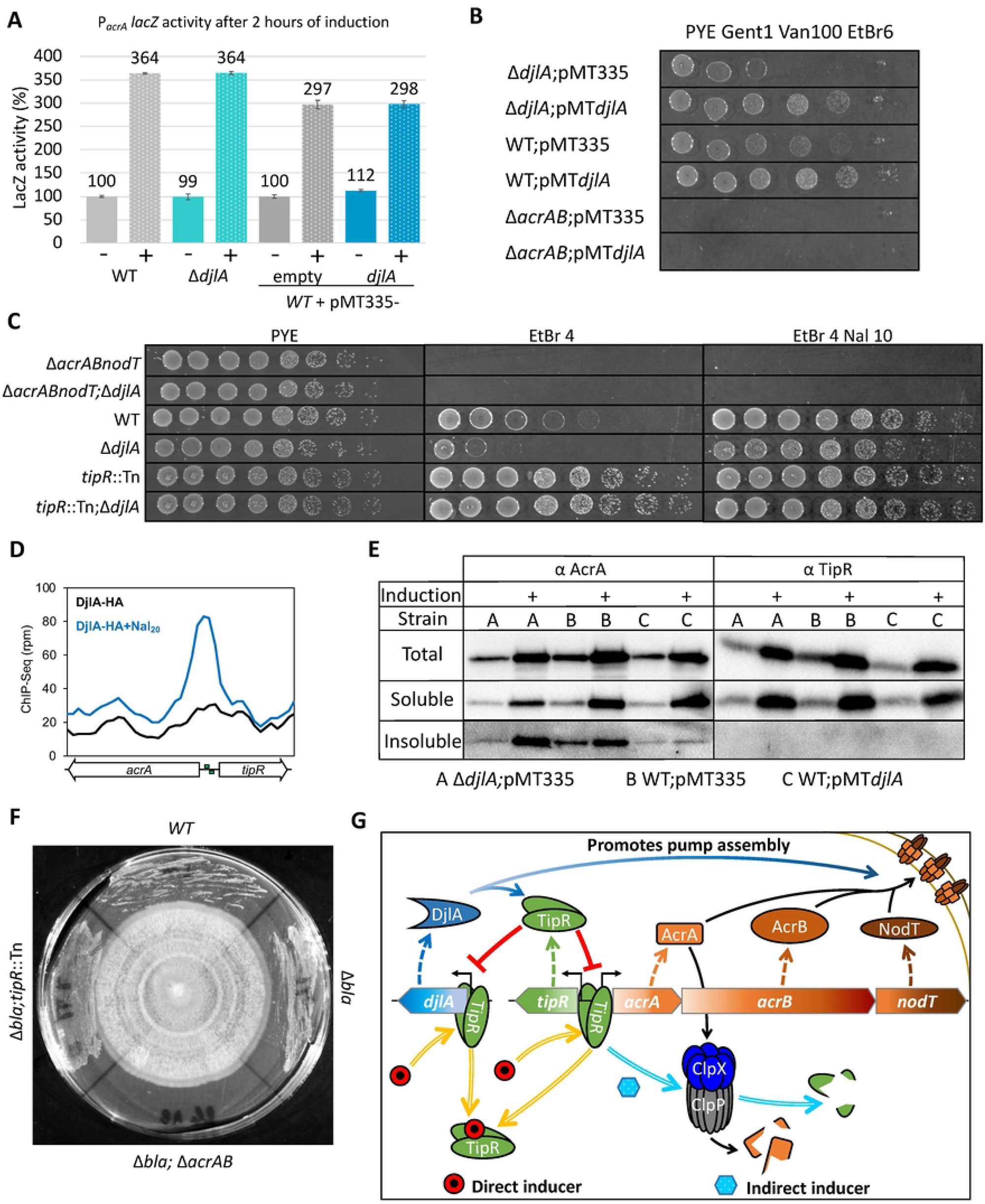
Role of DjlA co-chaperone in the TipR-dependent control loop. A) Measurement of β-galactosidase expressed from P_*acrA*_-*lacZ* in different *C. crescentus* Δ*djlA* mutants before or after a 2 hour induction with nalidixic acid 10 µg/mL (+) and/or vanillate 100 µM (V). All measurements are indicated in percentage of relative to the basal activity of the *WT* control before induction. B) Efficiency of plating (EOP) assay determined by ten-fold serial dilutions of Δ*djlA* and Δ*acrAB-nodT* mutants on plates containing ethidium bromide (EtBr) to probe for efflux pump activity. All strains contain pMT335 plasmid or a derivative (pMT335-*djlA*) expressing DjlA, grown on vanillate (Van) at 100 µM and gentamicin (Gent) 1 µg/mL. C) Efficiency of plating (EOP) assay by ten-fold serial dilutions of Δ*acrAB-nodT* and Δ*djlA* mutants on plates with ethidium bromide (EtBr in µg/mL) to probe for pump efficiency. Nal induction was performed at 10 µg/mL, vanillate (Van) at 100 µM and gentamicin (Gent) in µg/mL. D) ChIP-Seq analysis of DjlA-HA expressed from the *xylX* locus in *WT* cells before and after NAL induction. DjlA-HA was pulled down using an anti-HA affinity matrix. The *acrA*-*tipR* promoter region was the only region showing a > 2.9-fold increase in abundance compared to the control ChIP-seq performed with cells that were not treated with NAL for 30 minutes. E) Immunoblot analysis of aggregated cellular proteins in various strains A = Δ*djlA*, B = *WT* and C = *WT* pMT*djlA*. Inductions (+) were performed for 2 hours with 10 μg/mL of nalidixic acid (Nal). The total fraction contains the cell lysate. The insoluble fraction corresponds to the material that remained in the pellet after several extractions with 1% Triton X-100. Strain C was grown in PYE with gentamicin 1 µg/mL and vanillate 100µM. F) AcrAB-NodT promotes growth of C. crescentus in the vicinity of an unknown fungus. *C. crescentus* strains grown in contact with an unknown fungus on a PYE plate. The fungus was pre-grown the plate for 2 weeks, after which the C. crescentus strains were streaked from the edge of the plate towards the center until in contact with the fungus. The plate was then incubated at 30°C for another 72 hours. The fungus had been isolated in the laboratory as a contaminant on a PYE plate. AcrAB-NodT encoded in the *C. crescentus* chromosome protects against (a) compound(s) produced by the fungus to enable growth of *C. crescentus* in the vicinity of the fungus. G) Proposed model of the *acrAB-nodT* operon and *djlA* gene regulation, induction and function.

Since co-chaperones like DjlA typically assist in protein folding and/or disaggregation under stress conditions, we reasoned that NAL-induced co-expression of DjlA with AcrAB-NodT may serve to manage a stress condition (for the envelope) imposed by the massive synthesis and assembly of the AcrAB-NodT structure, perhaps facilitating disaggregation of AcrA before secretion from the cytoplasm where DjlA is located. In support of this idea, pull-down experiments (Figure S19) using a DjlA variant with a C-terminal HA-tag (DjlA-HA) revealed that DjlA directly or indirectly associates with AcrA and TipR. Moreover, ChIP-Seq experiments (Figure 6D) indicate that DjlA-HA can associate with TipR at the P_*acrA*_ chromosomal site *in vivo* following exposure to NAL. In fact, we noted a near 3-fold increase in P_*acrA*_ abundance when DjlA-HA was immunoprecipitated from NAL-treated cells *versus* untreated cells, suggesting that i) DjlA-HA binds P_*acrA*_ via TipR and that ii) P_*acrA*_ occupancy by DjlA-HA is enhanced by NAL.

Next, we explored whether DjlA also influences aggregation of TipR or AcrA. To test this, we first induced *WT* cells, Δ*djlA* cells and DjlA-overexpressing cells (*WT*+ pMT335-*djlA*) with NAL and then separated extracts into soluble and (Triton X-100) insoluble fractions by centrifugation (Figure 6E). Immunoblotting revealed AcrA to be more abundant in the soluble fraction from *WT* cells *versus* those from Δ*djlA* cells. Moreover, little aggregated AcrA was detected in the insoluble fraction from *WT* cells overexpressing DjlA, versus those from *WT* or Δ*djlA* cells. By contrast, no major difference in TipR abundance was detectable in these fractions.

To determine if DjlA also acts on other (envelope) proteins, we conducted LC-MS/MS (liquid chromatography followed by tandem mass spectrometry) analyses of the DjlA-HA pull-downs described above (Figure S19). Pull-downs of DjlA-HA expressing cells grown in the absence of NAL, revealed an impressive number of <u>T</u>on<u>B</u>-<u>d</u>ependent receptors (TBDR) and porins destined for the OM as clients of DjlA. In the presence of NAL, there was a drastic reduction of these number of OM proteins (OMPs) with a concomitant increase in abundance of AcrA, AcrB and Nod, suggesting that DjlA has higher affinity for efflux pump components than for OMPs which is consistent with its role in enhancing efflux activity by preventing its aggregation.

## Discussion

Here we elucidated an integrated homeostatic loop (Figure 6G) controlling the transient induction of the AcrAB-NodT efflux pump in *C. crescentus*, reinforced at the post-translational level by the co-induced DjlA co-chaperone. Adaptive resistance to β-lactam antibiotics is conferred by the inducible AcrAB-NodT efflux pump whose overexpression from TipR-repressed promoter can be triggered by different classes of chemicals: some are antibiotics, while others are general noxious molecules. Remarkably, we unravelled two mechanisms of control acting through TipR, with one class of compounds interfering with DNA-binding of TipR, while others act on TipR stability. Importantly, we uncovered corresponding mutations in TipR that recapitulate these two divergent induction mechanisms. Our deep mutational scanning using forward selection for β-lactam resistance mapped three major domains in the TipR primary structure (g, h, and i in Figure 2A): the N-terminal DNA-binding, the central domain and the extreme C-terminus for (de)stabilization. Mutations in each of these domains can lead to de-repression of P_*acrA*_ with overexpression of AcrAB-NodT and cephalosporin resistance. Additionally, we used a two-step antibiotic selection regime and Mut-Seq to show that a second (elevated) level of resistance can still be acquired in *tipR* mutant cells that overexpress AcrAB-NodT by with missense mutations in AcrB, augmenting cephalothin resistance to 3-orders or magnitude compared to regular AcrAB-NodT overexpression strains.

Although inducible overexpression of AcrAB-NodT is critical for survival against certain drugs or toxic molecules (also against natural molecules, Figure 6F), it also comes at a cost for envelope integrity and (cell shape) homeostasis. We discovered that long-term overexpression of the AcrAB-NodT envelope-spanning system results in strong OM perturbations that can sensitize cells towards antibiotics such as the cell wall (CW)-targeting antibiotic vancomycin that is normally efficiently excluded from cells by the barrier function of the OM. This barrier function is apparently compromised by excessive and/or long-term efflux pump (activity), perhaps owing to the OM detachment from other envelope layers membrane blebs and periplasmic vesicles that occur. It stands to reason that these defects should transiently result in a discontinuous OM or openings through which drugs like vancomycin can subsequently enter. Thus, to minimize envelope integrity defects caused by long-term expression of AcrAB-NodT, cells must ensure that its expression is transient and that the repressed ground state can re-establish as quickly as possible after induction.

In addition to binding AcrAB-NodT, the cytoplasmic co-chaperone binds several TBDRs and other OMPs (see Figure S19). We found that upon AcrAB-NodT induction, DjlA is drawn to the preferred AcrAB-NodT substrate, in favour over other OMPs. Such a substrate preference switch may also accentuate the OM problems arising from extended AcrAB-NodT overexpression, perhaps leading to a reduction of OMPs that can tether the OM to the CW. However, since Δ*djlA* cells do not show these OM defects in the absence of induction, AcrAB-NodT induction on its own may suffice to cause OM detachment, perhaps by constraining space in the OM that would otherwise be available for the insertion such CW-binding OMPs in regions occupied by AcrAB-NodT when it is overexpressed. It is also possible that AcrAB-NodT pumps out a molecule needed for tethering the OM to the cell wall or creates an imbalance in the availability of OM components versus those needed for the cytoplasmic membrane.

Our finding that the TipR expression itself is also induced by NAL, along with AcrAB-NodT and DjlA, is consistent with the view that this joint genetic control sets the stage for re-establishing repression as soon as the inducing compound is removed from the cytoplasm (Figure 6G). While repression by TipR will then terminate transcription of *acrAB-nodT*, the cell must also reduce the levels of the induced AcrAB-NodT protein by proteolysis, rather than by dilution due to cellular division which is substantially slower than proteolysis. Indeed, AcrA is substrate of the ClpXP protease (Bhat et al., 2013) and our findings provide further evidence for interactions between TipR, AcrA and the degradasome. The ATP-dependent protease ClpP restrict the passage to its proteolytic chamber only small, unfolded peptides, through narrow pores. To degrade bigger molecules, the proteasome needs ClpX that recognize specific substrates, directly or using adaptors to unfold the targeted protein using ATP. This then allows the denatured polypeptide chain to be translocated to the ClpP inner chamber where it gets degraded. One explanation could be that quinolones recruit an adaptor to TipR, or the antibiotic molecules block proper folding of the repressor, rendering them recognisable by the ClpXP machinery.

In the absence of a known natural inducer, we sought and discovered 14, mostly synthetic molecules that trigger de-repression of TipR. Among the inducers described in this study, two molecules possess a curious behaviour: dopamine and aminolevulinic acid that do not induce AcrAB-NodT expression in short term. While aminolevulinic acid can be used as a natural heme precursor, dopamine is a catecholamine characterised by an iron binding domain. Both molecules are highly unstable and change conformation after time, in particular aminolevulinic acid that spontaneously dimerises into porphobilinogen, pseudoporphobilinogen or 2,5-dicarboxyethylpyrazine forming non-heme precursors (Tewari and Eggleston, 2018). As those molecules share structural similarities with efflux pump substrates, we speculate that aminolevulinic acid is not an inducer in this form, but its degradation or dimer products might trigger expression of the efflux pump, perhaps through an iron stress response. In support of this idea, our RNA-Seq analysis of NAL-treated cells shows an increase of expression in the transcript encoding the 5-aminolevulinic acid synthase, hinting at a purpose of *acrAB-nodT* expression control and inducers, with aminolevulinic acid derivatives activating the AcrAB-NodT expression to export an unusable product from the cell.

AcrAB-NodT likely export natural products and metabolites that enter from the outside as well. Chemical warfare between microbes typically involves a myriad of toxic molecules, sometimes targeting different compartments, and thus protection against many molecules at once can be signalled through a single inducer entering the cytoplasm. Indeed, a fungus isolated in the lab prevents growth of *C. crescentus* cells lacking AcrAB-NodT in proximity, but not cells expressing it (Figure 6F), thereby illustrating the importance of AcrAB-NodT in protecting bacteria towards natural toxic metabolites that likely also induce the system. However, we noted that the contribution of AcrAB-NodT-mediated efflux is also detectable without NAL-induction in assays of *WT* and for Δ*acrAB-nodT* cells using discs containing the cephalosporins cefepime or cefotaxime (Figure S1B). AcrAB-NodT also confers resistance to other antibiotics including erythromycin or other noxious compounds such as EtBr in the uninduced state, indicating that AcrAB-NodT also expels compounds other than β-lactams. β- lactams do not induce AcrAB-NodT expression, a circumstance that is likely explained by the fact that these drugs are not known to enter the cytoplasm, unlike smaller molecules like EtBr or NAL. While this property explains why resistance to β-lactams in *C. crescentus* is an adaptive trait, it may also confer multi-resistance to wide range of toxic molecules with a minimal range of inducers of the efflux pump.

Collectively, our findings explain why AcrAB-NodT control on at least two levels (TipR and DjlA) is critical for transient efflux activation that protects bacterial cells in the wild from noxious molecules, but also from antibiotic treatment in the clinical setting.

## MATERIALS AND METHODS

### Growth conditions

All the *C. crescentus* strains were cultivated in peptone-yeast extract (PYE) and incubated at 30°C. and ФCr30-mediated generalized transductions were done as described (Ely, 1991). *E. coli* strains were grown in Luria broth at 37°C. All media were supplemented with the appropriate antibiotics or indicated inducer. The strain used for cloning is *E. coli* EC100D while the strain used for protein expression is *E. coli* BL21(DE3). The list of strains is below. As both wild-type (*WT*, NA1000) and Δ*bla C. crescentus* cells are naturally resistant to Colistin (COL, 4 µg/mL) or aztreonam (AZT, 3 µg/mL), COL or AZT was used to counter-select *E. coli* Tn delivery strains for Tn (*himar1*) mutagenesis encoded on plasmid pHPV414 (Viollier et al., 2004). The *himar1* encodes resistance to kanamycin and therefore the PYE plates also contained kanamycin (20 µg/mL, KAN^20^) and CEF^10^ or PIR^40^.

### β-galactosidase assays

All the β -galactosidase assays were done using freshly electroporated strains harboring the pLac290 plasmid with transcriptional fusions between the studied promoters and *lacZ*. The assays were done at room temperature. Between 50 and 100 µL of cells (OD_600nm_= 0.3-0.7) were lysed in 30 µL of chloroform and vigorously mixed with Z buffer (60 mM Na2HPO4; 40 mM NaH2PO4; 10 mM KCl and 1 mM MgSO4; pH 7) to obtain a final volume of 800 µL. Followed by the addition of 200 µl of ONPG (Ortho-nitrophenyl-β-D-galactopyranoside, at 4 mg/mL in 0.1 M potassium phosphate, pH 7) to begin the reaction. Assays were stopped using 500 µL of 1 M Na2CO3 when the solution turned light yellow. The OD420nm of the supernatant was collected and use to calculate the Miller units as follows: U=(OD420*1000)/(OD_600_*t(min)*v(ml)). Error was calculated as standard deviation from at least three biological replicates.

### Kirby-Bauer disk diffusion susceptibility test

In a 14 cm diameter Petri dish, 50 mL of 1.5% agar media supplemented with indicated antibiotic and/or inducer were cast. After the polymerization, 12 mL of 0,375% agar media maintained at 50°C were mixed with 400 µL of overnight bacterial culture prior to be spread evenly on top of the solid media. Antibiotics disk were ordered from Bio-Rad or made in the lab using sterile disks. The pictures were taken after the plates were incubated overnight at appropriate temperate. Images were captured on a Bio-Rad illuminator and with the Image Lab 4.1 software.

#### Efficiency of plating

According to the OD_600_, the same quantity cells from an overnight culture were place in the first line of a 96 well plates. All the wells were filled with 180 µL of appropriate media except the first line that was adjusted to 200 µL. The dilutions were done with 20 µL of the first line of wells transferred to the following line using a multi pipette, this step was repeated until the 8 lanes were done. Finally, 5 µL of each dilution was dropped on a Petri dish with the solid media supplemented with the appropriate chemical(s).

#### Microscopy

2-4 µL of an exponential phase culture of *Caulobacter crescentus* grown in PYE supplemented with indicated chemicals were immobilized on a thin layer of 1% agarose pad. Contrast microscopy pictures were taken using a phase 100x objective with an oil interface (Zeiss, alpha plan achromatic 100x/1.46 oil phase 3) using an Axio Imager M2 microscope (Zeiss), with appropriate filter (Visitron Sys-tems GmbH) and a cooled CCD camera (Photometrics, CoolSNAP HQ2) controlled via the Metamorph software (Molecular Devices).

#### Electrophoretic mobility shift assay (EMSA)

A Cy5 labeled probes were generated by PCR with primers chemically modified with the fluorescent dye. The probes were purified by agarose gel electrophoresis. The reaction took place in a buffer containing: 40 mM Tris-HCl (pH 7.6), 60 mM KCl, and 0.1% glycerol, 240 µg of calf DNA, 800 µg of BSA and 200 ng of labeled probe. After the addition of the indicated quantity of purified protein, the sample were place 30 min in the dark at room temperature. After a pre-run of 30 min, the samples were loaded and migrated by electrophoresis in a nondenaturing 4% Tris-Bore-EDTA acrylamide (19 :1) gel. After migration the samples were observed in a Bio-Rad illuminator (Chemidoc MP) using the pre-installed parameters for Cy5 imaging, pictures were collected through Image Lab 4.1 software (Bio-Rad).

#### Chemical library screening

The Maybridge Chemical Library was delivered aliquoted into 96-well plates and dissolved in 50% DMSO. Petri dish of 22 cm containing PYE kanamycin 10 µg/mL were prepared using the Kirby-Bauer technique described above, with a NA1000 strain carrying the pLac290 plasmid harboring a fusion between the P_*acrA*_ and the *nptII* kanamycin resistance gene. Potential inducers were detected when a growth area occur at the position of a drop after 48 hours of incubation at 30°C.

#### Immunoblot analysis/chase

The appropriate strain was grown for 2 to 4 hours at 30°C under constant agitation up to OD_600nm_ of 0.4 to 0.6, then the inducer was added, and the cells were grown for 2 additional hours. For chase experiment, the strain was grown similarly up to OD_600nm_ of 0,3-0,5, then chloramphenicol 5 µg/mL was supplemented with vigorous mixing, immediately followed by the addition of mentioned inducer. Protein samples from exponentially growing cells were separated on a SDS–polyacrylamide (37.5:1) gel electrophoresis and blotted on 0.45µm pore PolyVinyliDenFluoride (PVDF) membranes (Immobilon-P from Sigma Aldrich). Membranes were blocked for 2 hours with 1x Tris-buffered saline (TBS) (50 mM Tris-HCl, 150 mM NaCl [pH 8]) that contain 0.1% Tween-20% and 8% powdered milk, followed by overnight incubation with the primary antibodies diluted in the same milk solution. The polyclonal antisera to AcrA (1:15000), TipR (1:5000) and CCNA_00164 (1:20000) were used. The detection of primary antibodies was done using HRP-conjugated donkey anti-rabbit antibody (Jackson ImmunoResearch) with Western Blotting Detection System (Immobilon from Milipore) and an imaging was performed in Bio-Rad illuminator (Chemidoc MP, Biorad).

#### Protein purification

TipR protein was expressed from pET21 in *E. coli* BL21(DE3)/pLysS. Cells were cultivated in LB at 37°C and induced by 1 mM of IPTG for 4 hours, up to an OD_600nm_=0.5. The bacteria were harvested at 8000 RPM for 30 minutes at 4°C. The pellet was re-suspended in 25 mL of buffer (TrisHCL 40mM pH 7.6, 50mM KCl) before to be sonicated in a water–ice bath (15 cycles of 30 seconds ON, 30 seconds OFF). After centrifugation at 5000g for 20 minutes at 4°C and filtration through 0.22 µm filters, the proteins contained in the supernatant were purified under native conditions successively using Q-sepharose and Heparin columns (HiTrap systems from Cytia) according to manufacturer manual. Protein concentration was determined using the Bradford quantification method. Proteins were stored at -80°C in 40mM Tris-HCl pH 7.6, 50 mM KCl and 10% glycerol.

#### Two-step resistance selection and Mut-seq

To isolate the Δ*bla tipR*::Tn derivatives that grow on CEF^40^ (Δ*bla tipR*::Tn *acrAB-nodT\**), Δ*bla tipR*::Tn cells were plated on CEF^40^, resistant mutant clones were pooled and ФCr30-lysates were prepared from this pool. The *tipR*::Tn allele was transduced into Δ*bla* cells and transductants were selected on plates with KAN^20^ and then tested for growth on plates with CEF^40^. The *acrAB-nodT* locus was sequenced in several such CEF^40^ -resistant clones.

The Δ*bla* strain was spread on PYE cephalothin 10 µg/mL and incubated at 30°C for 48 hours. Around 10 000 colonies were harvested and pooled prior to DNA extraction using Ready-Lyse Lysozyme (Epicentre Lucigen) and DNAzol (ThermoFischer). PCR were performed using the Q5 High-Fidelity DNA Polymerase according to manufacturer instructions with a maximum number of 15 cycles. The amplicons were purified with GeneJET Gel Extraction Kit (ThermoFischer) and send to sequencing at the iGE3 genomic platform in CMU (University of Geneva – Switzerland). The mixed amplicons were sent to the Genomic platform iGE3 at the university of Geneva. Library preparation and sequencing were done using a HiSeq 2500 with 50-bp paired-end reads. Data analysis was done using Burrows-Wheeler Alignment Tool version 0.7.5a and Samtools version 1.2 prior to be blast against the *C. crescentus* NA1000 reference genome (NC_011916.1). SNPs with an abundance lower than 5% were considered background (PCR amplification errors) and excluded from the analysis.

#### RNA extraction, deep-sequencing and bioinformatics analysis

Overnight cultures in PYE of *C. crescentus* NA1000 (*WT*) were freshly restarted in 10ml of PYE (starting O.D.660nm∼0.05) and incubated at 30°C under agitation to reach an O.D.660nm∼0,5. Nalidixic acid treated cultures were supplemented with 20 μg/mL of antibiotic 30 minutes before being collected. First, to provide immediate stabilization of RNA, each cell cultures (4ml) are treated with 2 volumes of RNA Protect Bacteria reagent (Qiagen, Switzerland) according to the manufacturer’s protocol. Then, cells were lysed in a Ready-Lyse lysozyme solution (Epicentre Technologies) according to manufacturer’s instructions and lysates were homogenized through QiaShredder columns (Qiagen, Switzerland). The RNeasy Mini Kit (Qiagen, Switzerland) was used for total RNA extraction according to the manufacturer’s protocol which included a first DNase treatment using the On-column DNase I digestion kit (Qiagen, Switzerland). Extracted total RNA was subjected to a second DNase treatment using Promega RQ1 at 1 unit/μg of RNA. Another total RNA cleanup was performed after the DNase treatment. The RNA concentration was measured using a Nanodrop 1000 spectrophotometer (ThermoScientific, USA) and RNA quality was assessed using a 2100 Bioanalyzer Instrument (Agilent Technologies, USA). Two independent biological replicates were analysed per condition.

RNA-Seq library preparation and sequencing was performed at Fasteris SA (Geneva, Switzerland). Bacterial rRNA were removed from each total RNA samples and RNA-Seq libraries were prepared using the Ovation Complete Prokaryotic RNA-Seq library system kit according to the manufacturer’s instructions. Single-end runs were performed on an Illumina NextSeq 500 instrument (50 cycles), yielding several million reads (stored as fastq files). Using the web-based analysis platform Galaxy (https://usegalaxy.org), the single-end sequenced reads quality was checked (FastQC, Galaxy Version 0.72), and reads were mapped (Bowtie2, Galaxy Version 2.3.4.3) to the *C. crescentus* NA1000 genome (NC_011916.1). The tables of counts with the number of reads mapping to each gene feature were prepared using the htseq-count software, Galaxy Version 0.9.1. For each experimental series, the counts normalization and the statistical differential expression analysis was performed using the DESeq2 software, Galaxy Version 2.11.40.6. Sequence data have been deposited to the Gene Expression Omnibus (GEO) database (GSEXXXXXX accession, samples nos. GSMXXXXXXX– GSMXXXXXXX).

#### Chromatin Immuno Precipitation coupled to deep Sequencing (ChIP-Seq) and data analysis

Overnight cultures in PYE of *C. crescentus* NA1000(WT) or Δ*bla;* P*_xylX_*::P*_xyl_-djlA-HA* were freshly restarted in 80ml of PYE (starting O.D.660nm∼0.05) and incubated at 30°C under agitation with 0.3% xylose when necessary. Nalidixic acid treated culture was supplemented with 20 μg/mL of antibiotic 30 minutes before being fixed. Cultures of exponentially growing cells (O.D.660nm of 0.5) were supplemented with 10 μM sodium phosphate buffer (pH 7.6) and then treated with formaldehyde (1% final concentration) at RT for 10 minutes to achieve crosslinking. Subsequently, the cultures were incubated for an additional 30 minutes on ice and washed three times in phosphate buffered saline (PBS, pH 7.4). The resulting cell pellets were stored at -80°C. After resuspension of the cells in TES buffer (10 mM Tris-HCl pH 7.5, 1 mM EDTA, 100 mM NaCl) containing 10 mM of DTT, the cell resuspensions were incubated in the presence of Ready-Lyse lysozyme solution (Epicentre, Madison, WI) for 10 minutes at 37°C, according to the manufacturer’s instructions. Lysates were sonicated (Bioruptor® Pico) at 4°C using 15 bursts of 30 seconds to shear DNA fragments to an average length of 0.2-0.5 kbp and cleared by centrifugation at 14,000 rpm for 2 min at 4°C. The volume of the lysates was then adjusted (relative to the protein concentration) to 1 mL using ChIP buffer (0.01% SDS, 1.1% Triton X-84 100, 1.2 mM EDTA, 16.7 mM Tris-HCl [pH 8.1], 167 mM NaCl) containing protease inhibitors (Roche) and pre-cleared with 80 μl of Protein-A agarose (Roche, www.roche.com) and 100 μg BSA. 5% of each pre-cleared lysate were reserved as total input samples (negative control samples). The pre-cleared lysates were then incubated overnight at 4°C with polyclonal rabbit anti-TipR antibodies (1:400 dilution) (Kirkpatrick and Viollier, 2014) or monoclonal rabbit anti-HA antibodies (1:250 dilution) (Clone 114-2C-7, Merck Millipore).. The immuno-complexes were captured after incubation with Protein-A agarose beads (pre-saturated with BSA) during a 4 h incubation at 4°C and then, washed subsequently with low salt washing buffer (0.1% SDS, 1% Triton X-100, 2 mM EDTA, 20 mM Tris-HCl pH 8.1, 150 mM NaCl), with high salt washing buffer (0.1% SDS, 1% Triton X-100, 2 mM EDTA, 20 mM Tris-HCl pH 8.1, 500 mM NaCl), with LiCl washing buffer (0.25 M LiCl, 1% NP-40, 1% deoxycholate, 1 mM EDTA, 10 mM Tris-HCl pH 8.1) and finally twice with TE buffer (10 mM Tris-HCl pH 8.1, 1 mM EDTA). The immuno-complexes were eluted from the Protein-A agarose beads with two times 250 μL elution buffer (SDS 1%, 0.1 M NaHCO3, freshly prepared) and then, just like total input samples, incubated overnight with 300 mM NaCl at 65°C to reverse the crosslinks. The samples were then treated with 2 μg of Proteinase K for 2 hours at 45°C in 40 mM EDTA and 40 mM Tris-HCl (pH 6.5). DNA was extracted using phenol:chloroform:isoamyl alcohol (25:24:1), ethanol-precipitated using 20 μg of glycogen as a carrier and resuspended in 50 μL of DNAse/RNAse free water.

Immunoprecipitated chromatin was used to prepare sample libraries used for deep sequencing at Fasteris SA (Geneva, Switzerland). ChIP-Seq libraries were prepared using the DNA Sample Prep Kit (Illumina) following manufacturer instructions. Single-end run was performed on an Illumina Next-Generation DNA sequencing instruments (NextSeq High), 50 cycles were performed and yielded several million reads per sequenced samples. The single-end sequence reads stored in FastQ files were mapped against the genome of C. crescentus NA1000 (NC_011916.1) using Bowtie2 version 2.4.2.+galaxy0 available on the web-based analysis platform Galaxy (https://usegalaxy.org) to generate the standard genomic position format files (BAM). ChIP-Seq reads sequencing and alignment statistics are summarized in Table S3. Then, BAM files were imported into SeqMonk version 1.47.2 (http://www.bioinformatics.babraham.ac.uk/projects/seqmonk/) to build ChIP-Seq normalized sequence read profiles. Briefly, the genome was subdivided into 50 bp, and for every probe, we calculated the number of reads per probe as a function of the total number of reads (per million, using the Read Count Quantitation option). Analyzed data illustrated in Figure 1F and 6D are provided in Table S3. Using the web-based analysis platform Galaxy (https://usegalaxy.org), TipR ChIP-Seq peaks were called using MACS2 Version 2.1.1.20160309.6 (No broad regions option) relative to the total input DNA samples. The q-value (false discovery rate, FDR) cut-off for called peaks was 0.05. Peaks were rank-ordered according to their fold-enrichment values (Table S3, Peaks with a fold-enrichment values >4 for TipR were retained for further analysis). Consensus sequences common to the 4 enriched TipR-associated loci were identified by scanning peak sequences (+ or - 75 bp relative to the coordinates of the peak summit) for conserved motifs using MEME (http://meme-suite.org/) (Bailey et al., 2009a; Bailey et al., 2009b). Sequence data have been deposited to the Gene Expression Omnibus (GEO) database (XXXXXX series, accession nos. XXXXXX–XXXXXX).

### Plunge freezing of *Caulobacter crescentus* cells

*C. crescentus* cells were mixed with 10 nm Protein A conjugated colloidal gold particles (1:10 v/v, Cytodiagnostics) and 4 µl of the mixture was applied to a glow-discharged holey-carbon copper EM grid (R2/1 or R2/2, Quantifoil). The grid was automatically backside blotted for 4-6s in a Mark IV Vitrobot (Thermo Fischer Scientific) by using a Teflon sheet on the front pad, and plunge-frozen in a liquid ethane-propane mixture (37%/63%) cooled by a liquid nitrogen bath. Frozen grids were stored in liquid nitrogen until loaded into the microscope.

### Cryo-electron tomography

*C crescentus* cells were imaged by cryo-electron tomography (cryoET)(Weiss et al., 2017). Image were recorded on Titan Krios 300 kV microscopes (Thermo Fisher Scientific) equipped with a Quantum LS imaging filter operated at a 20 eV slit width and K2 or K3 Summit direct electron detectors (Gatan). Tilt series were collected using a bidirectional tilt-scheme from -60 to +60° in 2° increments. Total dose was 130-150 e^-^/Å and defocus was kept at -8 µm. Tilt series were acquired using SerialEM (Mastronarde, 2005), drift-corrected using alignframes, reconstructed and segmented using the IMOD program suite(Kremer et al., 1996). To enhance contrast, tomograms were deconvolved with a Wiener-like filter(Tegunov and Cramer, 2019).

### *In vivo* protein aggregation assay

40 mL of exponentially growing cells (OD_600_ between 0,4 and 0,6) were rapidly cooled in ice bath and pelleted by centrifugation (6000g, 10 minutes). All steps were performed at 4°C and buffers were all supplemented with protease inhibitor cocktail from Roche (cOmplete tablets, EASYpack). The pellets were washed once in buffer A (50 mM Tris/HCl pH8.0, 150 mM NaCl) and resuspended in 300 µL of buffer A supplemented with 100 U/mL Ready-Lyse Lysozyme (Epicentre Lucigen), 10 µg/mL of DNAse I (Roche) and 10 µg/mL of RNAse A (Invitrogen). Cells were lysed in a Bioruptor (Diagenode) (set to high, 15 cycles for 30 seconds at 4°C). Lysates were centrifuged (5000g, 10 minutes) two times to remove non-lysed cells. The protein concentration of the lysate was quantified by Bradford assay and an aliquot was harvested as the Total fraction. The remaining samples were centrifuged (14000g, 30 minutes) to pellet the insoluble protein fraction. The supernatant was kept as the Soluble fraction. The pellets were then resuspended in 300 μL buffer A supplemented with 1% (v/v) Triton X-100 with and incubated for 1 hour on ice with regular vortexing prior to the sonication in a Bioruptor (set to high, 1 cycle for 30 seconds) followed by a centrifugation (20000g, 20 minutes) for washing. This procedure was repeated three times. The protein pellet was resuspended in 100 μL 1xSDS loading buffer prior heating to 95°C for 10 minutes.

### Immunoprecipitation

50 mL of exponentially growing cells were harvested by centrifugation (20 minutes, 4500 rpm at 4°C). The pellets were washed in 50 mL 1xPBS and centrifuged for 15 minutes with 6000rpm at 4°C, then washed once in 1 mL 1xPBS before centrifugation (for 5 minutes, 14000 rpm, at 4°C). The pellets were resuspended in 1 mL TES (10 mM Tris-HCl pH 7.5; 1 mM EDTA; 100 mM NaCl) containing protease inhibitor (cOmplete tablets, EASYpack from Roche) at 2 tabs per 50 mL and 100 U of Ready-lyse/ml of culture (Ready-Lyse Lysozyme from Epicentre Lucigen) prior 10 minutes of incubation at room temperature. Each sample were supplemented with 50 μL NP-40 10% (AppliChem); 1 μL EDTA 0.5 M pH 8.0 (Sigma); 10 μL MgCl2 1 M; 10 μL DNAse I (Roche) 10 mg/mL; 10 μL RNAse A (Invitrogen) 10 mg/mL and incubated 20 minutes at room temperature with constant agitation. The non-lysed cells were removed by centrifugation (10 minutes, 8000 rpm at room temperature). The supernatant was harvested and centrifuged for 20 minutes with 14000 rpm at 4°C. The remaining supernatant was kept as IP input.

For each sample, 25 μL of GFP-trap (GFP-Trap_A from Chromotek) or anti-HA affinity matrix beads (Roche) was rinsed four times with 1 mL TES buffer and centrifugation 2 minutes at 3000 rpm. Then, 25 µL of beads were added to each IP input and left overnight at 4°C with agitation. The beads were harvested by centrifugation (2 min at 3000 rpm) and washed 4 times with Wash buffer (10 mM Tris-HCl 7.5; 150 mM NaCl; 0.5 mM EDTA; 0.5 % n-dodecyl-β-D-maltoside (DDM)). The beads were then resuspended in 40 μL of Laemmli sample buffer 2x (125 mM Tris-HCl pH6.8 ; 4% SDS ; 20% glycerol ; 10% B-mercaptoethanol ; 0.004% bromophenol blue) and incubated 10 minutes at 95°C. Each sample were centrifugated for 2 minutes at 3000 rpm and the supernatant was collected (without pipetting the beads). Five or ten μL of each collected fraction was analysed by SDS PAGE and Coomassie or immunoblotting.

## Acknowledgements

This work was supported by a Swiss National Science Foundation (Grant CRSII5_198737 to P.H.V), the Canton de Genève. We acknowledge instrument access at the imaging platform ScopeM at ETH Zürich. Work in Zürich was supported by the NOMIS foundation J.C. owes special thanks to the Fondation Ernst et Lucie Schmidheiny and wishes to acknowledge Sabine Quindou, Jamy Gourmaud and Frédéric Courant for nurturing an entire generation’s curiosity..

## Data availability

Representative reconstructed tomograms (EMD-XXXXX, EMD-XXXXX, EMD-XXXXX, EMD-XXXXX, EMD-XXXXX) have been deposited in the Electron Microscopy Data Bank.

## Strains, plasmids and primers used in this study

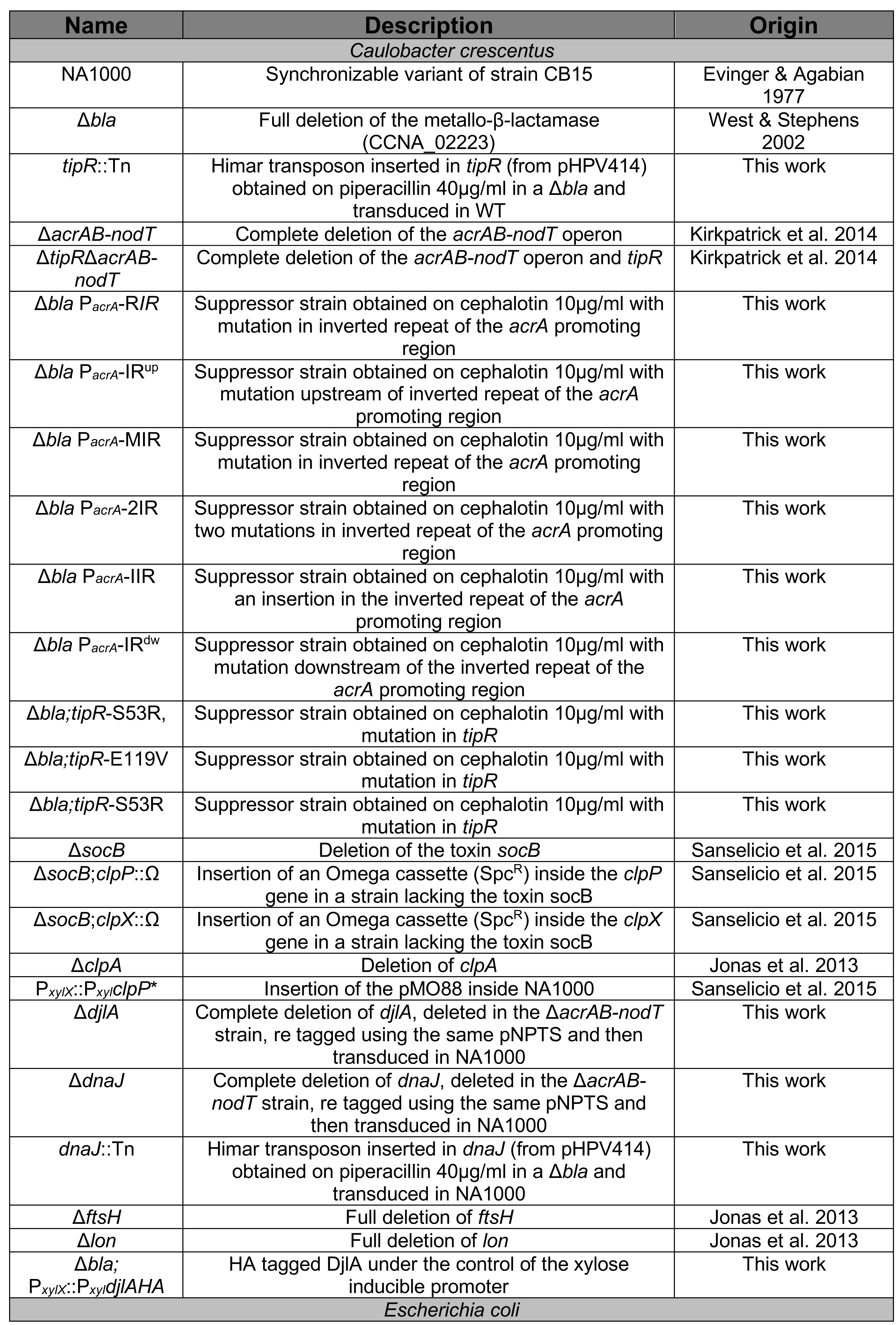
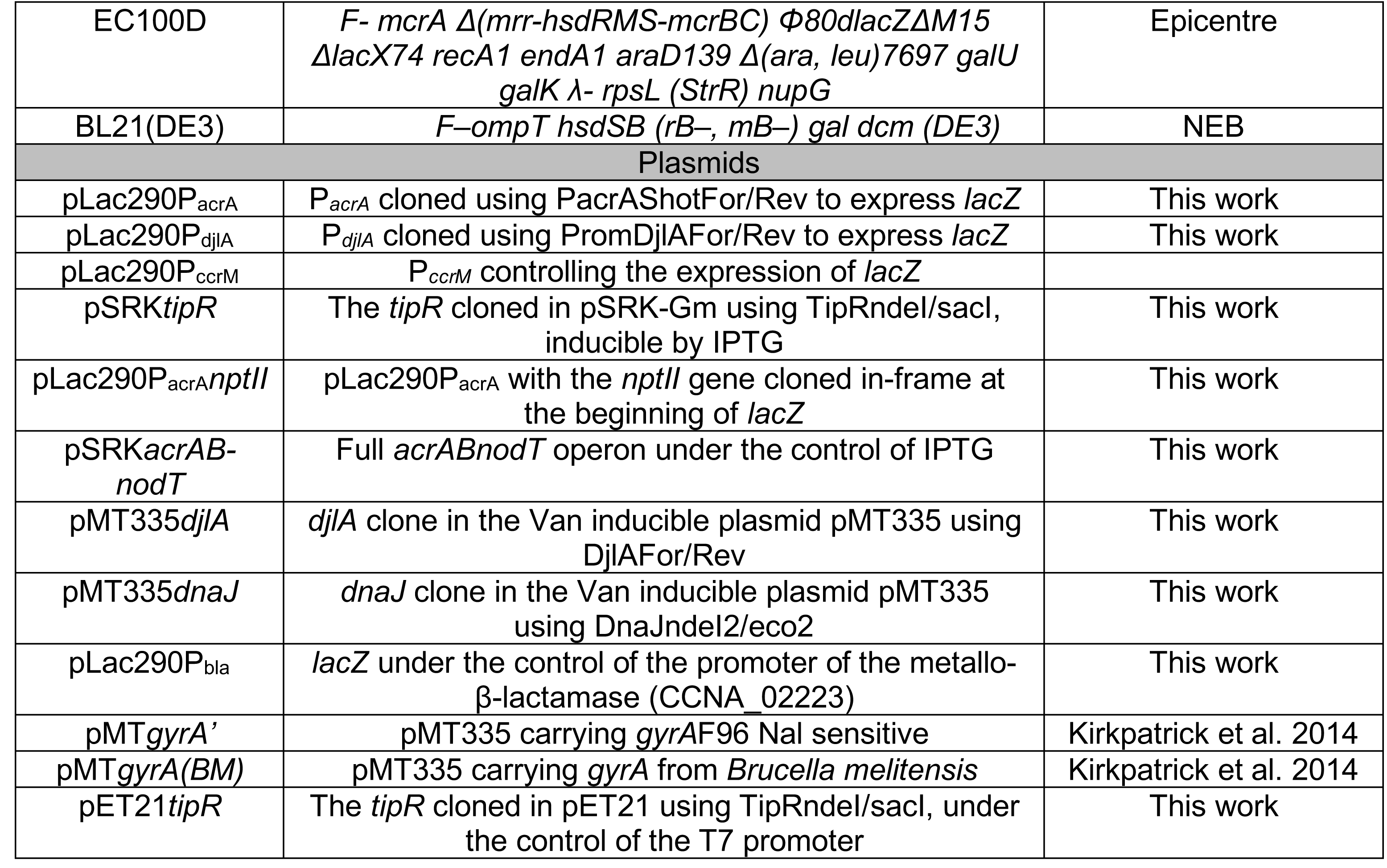
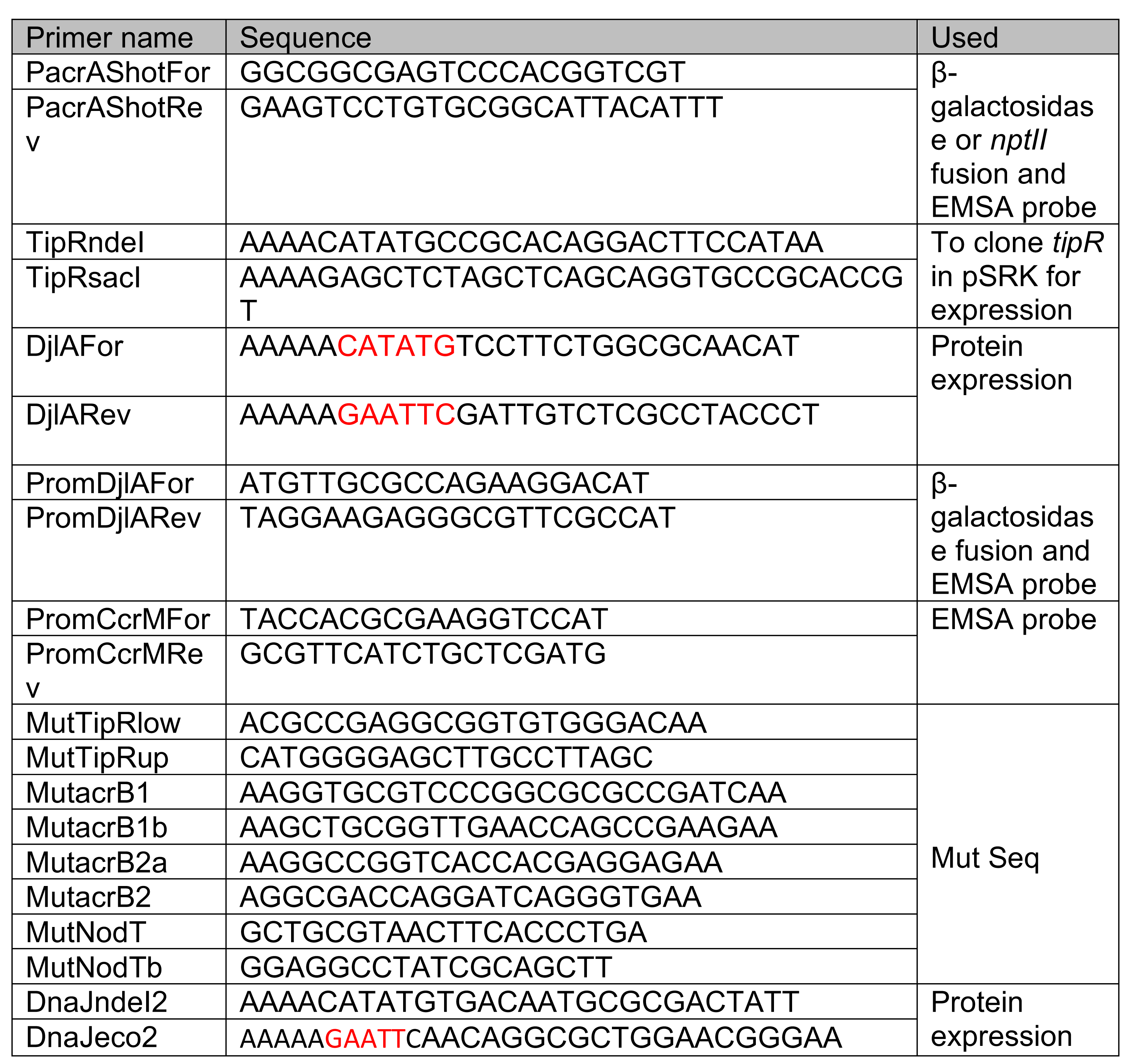

## Legends to supplementary figures

**Figure S1:**
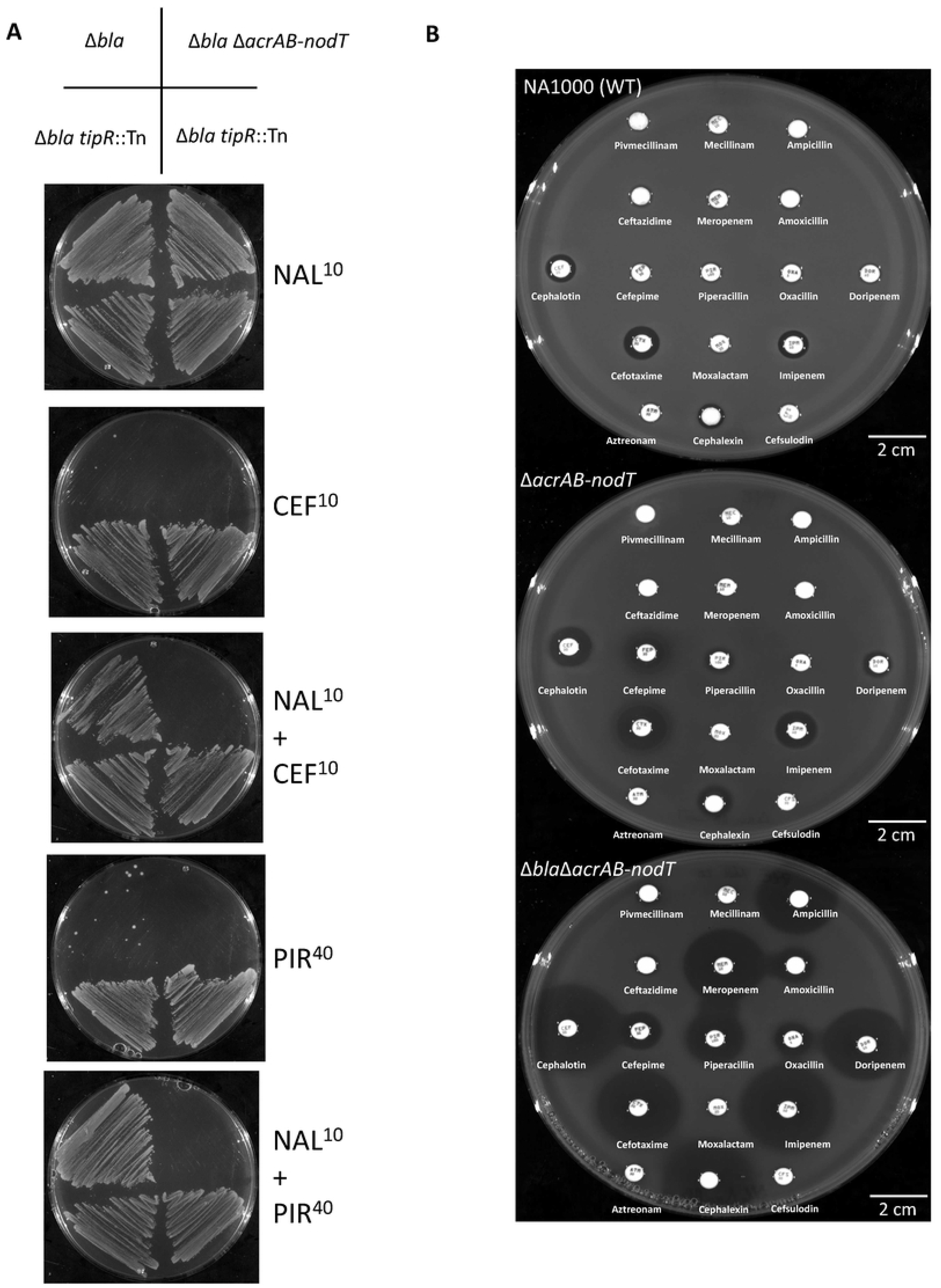
Resistance of *tipR* mutants to β-lactams. A) Growth of *C. crescentus* mutant strains on plates containing NAL^10^ (nalidixic acid, 10 µg/mL), CEF^10^ (cephalothin 10 µg/mL) and/or PIR^40^ (piperacillin 40 µg/mL) for 3 days. B) Antibiograms of *C. crescentus* strains on PYE. Antibiotic discs, from top left to bottom right: Pivmecillinam 20 µg, Mecillinam 10 µg, Ampicillin 100 µg, Ceftazidime 40 µg, Meropenem 10 µg, Amoxicillin 4 µg, Cephalothin 30 µg, Cefepime 30 µg, Piperacillin 100 µg, Oxacillin 5 µg, Doripenem 10 µg, Cefotaxime 30 µg, Moxalactam 30 µg, Imipenem 10 µg, Aztreonam 30 µg, Cephalexin 40 µg, Cefsulodin 30 µg.

**Figure S2:**
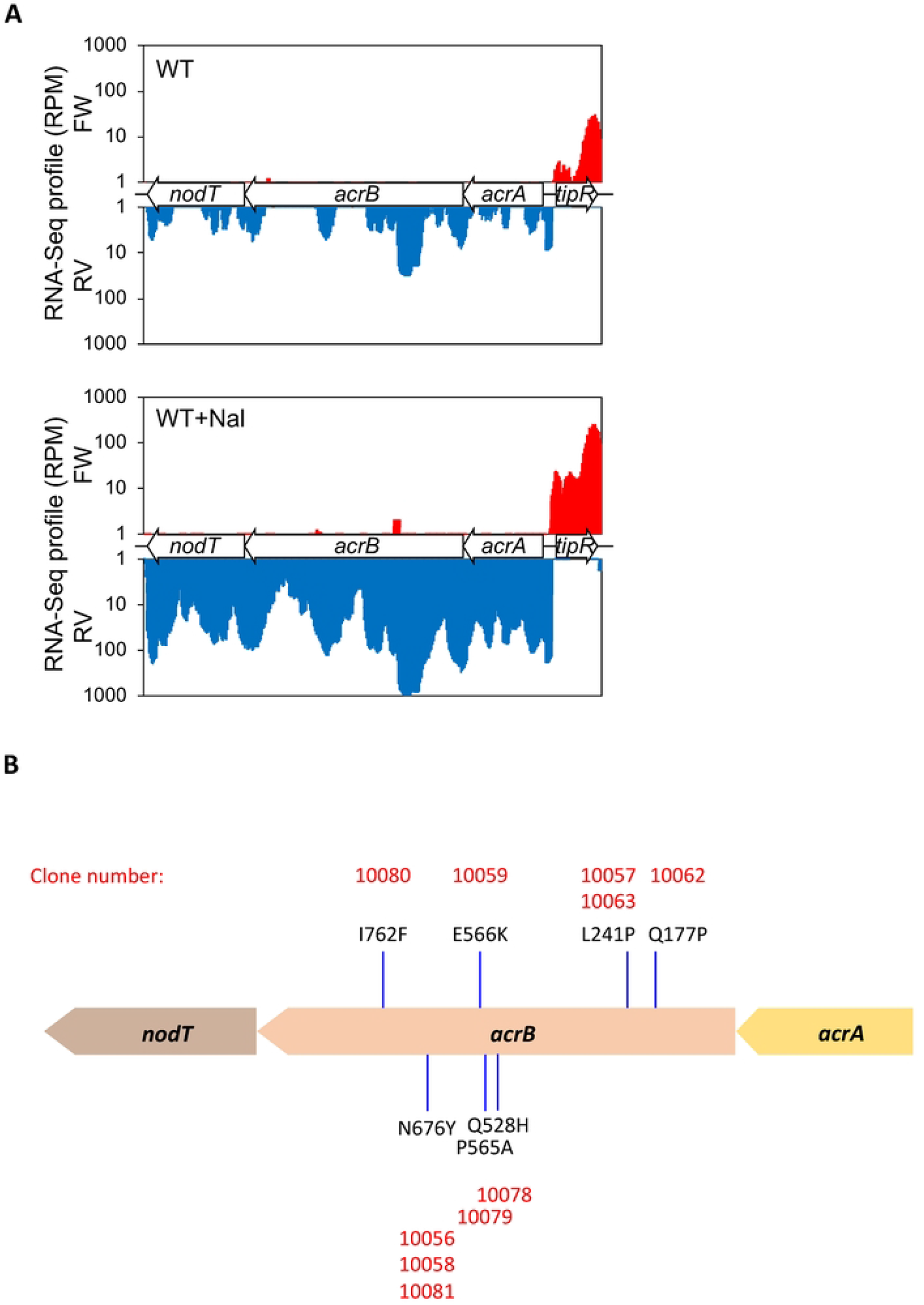
RNA-Seq analysis of the *tipR-acrAB-nodT* locus. A) Representation of the reads (represented as Reads Per Million (RPM)) obtained from the RNA-Seq experiment covering the *tipR* and *acrAB-nodT* region in the NA1000 strain (WT). Induction was performed on exponentially grown cells in PYE for 30 minutes with nalidixic acid (Nal) 20µg/mL. Red curves represent the reads in forward orientation (FW), blue curves represent the reads in reverse orientation (RV). B) Scheme showing the location of the AcrB mutations conferring high level of cephalothin resistance (CEF^40^) to Δ*bla tipR::*Tn cells.

**Figure S3:**
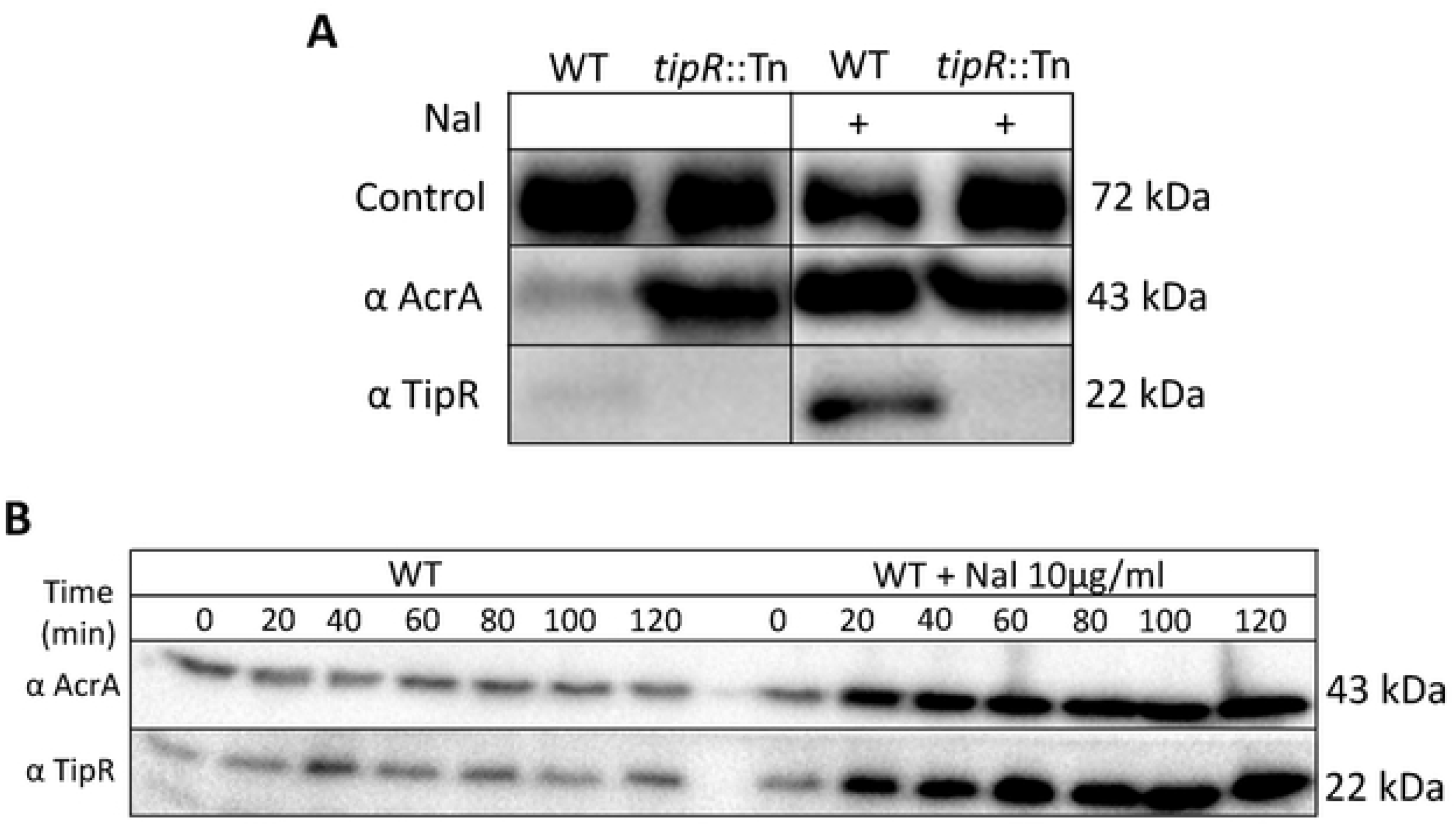
Induction of AcrA and TipR by NAL. A) Immunoblot conducted with anti-AcrA and anti-TipR on NA1000 (*WT*) and *tipR* mutant (*tipR*::Tn) cells exponentially grown in PYE before and after 2 hours of induction of Nal 20 µg/mL. Loading control represents CCNA_00163 revealed with antibodies to CCNA_00163. B) Immunoblot conducted with anti-AcrA and anti-TipR during a time course experiment on NA1000 (*WT*) for 2 hours in the presence and absence of alidixic acid (Nal) 10 µg/mL in PYE.

**Figure S4:**
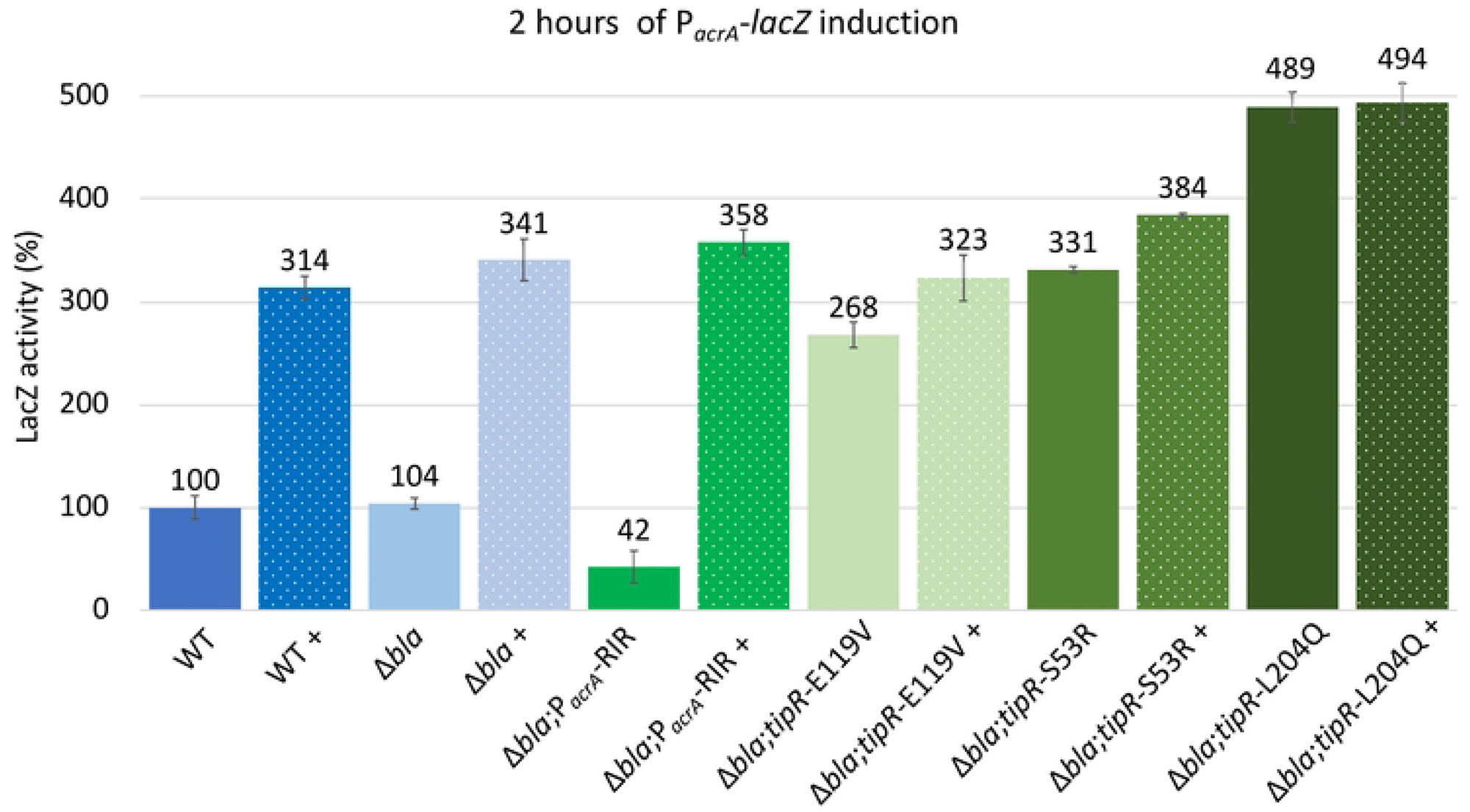
P*a*crA activity in various *tipR* mutants. β-galactosidase activity expressed from P_*acrA*_-*lacZ* in various mutants. Induction (+) was for 2 hours with nalidixic acid (Nal, 10 µg/mL). All levels are indicated in percentage of expression regarding the basal level of the *WT* (NA1000) without induction. All strains carry additionally the pP_*acrA*_-*lacZ* promoter probe plasmid.

**Figure S5:**
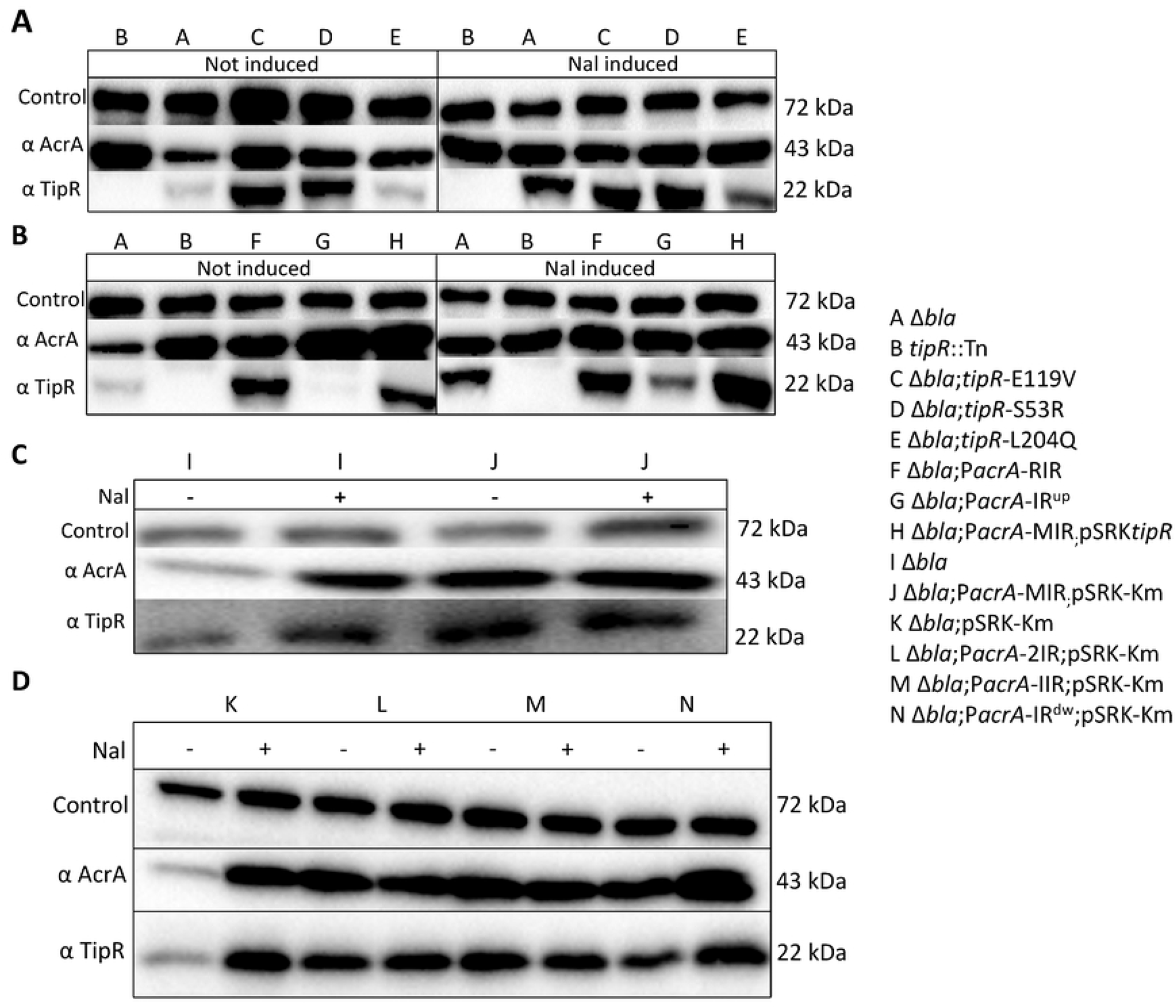
Abundance of TipR and AcrA in various *tipR* and PacrA point mutants. Immunoblots of different P_*acrA*_ (B, C and D) or *tipR* (A) mutants. All inductions (+) were performed for 2 hours at 10 µg/mL of Nalidixic acid (Nal) on exponentially grown cells in PYE. Blots were also probed with antibodies to CCNA_00163 as loading control.

**Figure S6:**
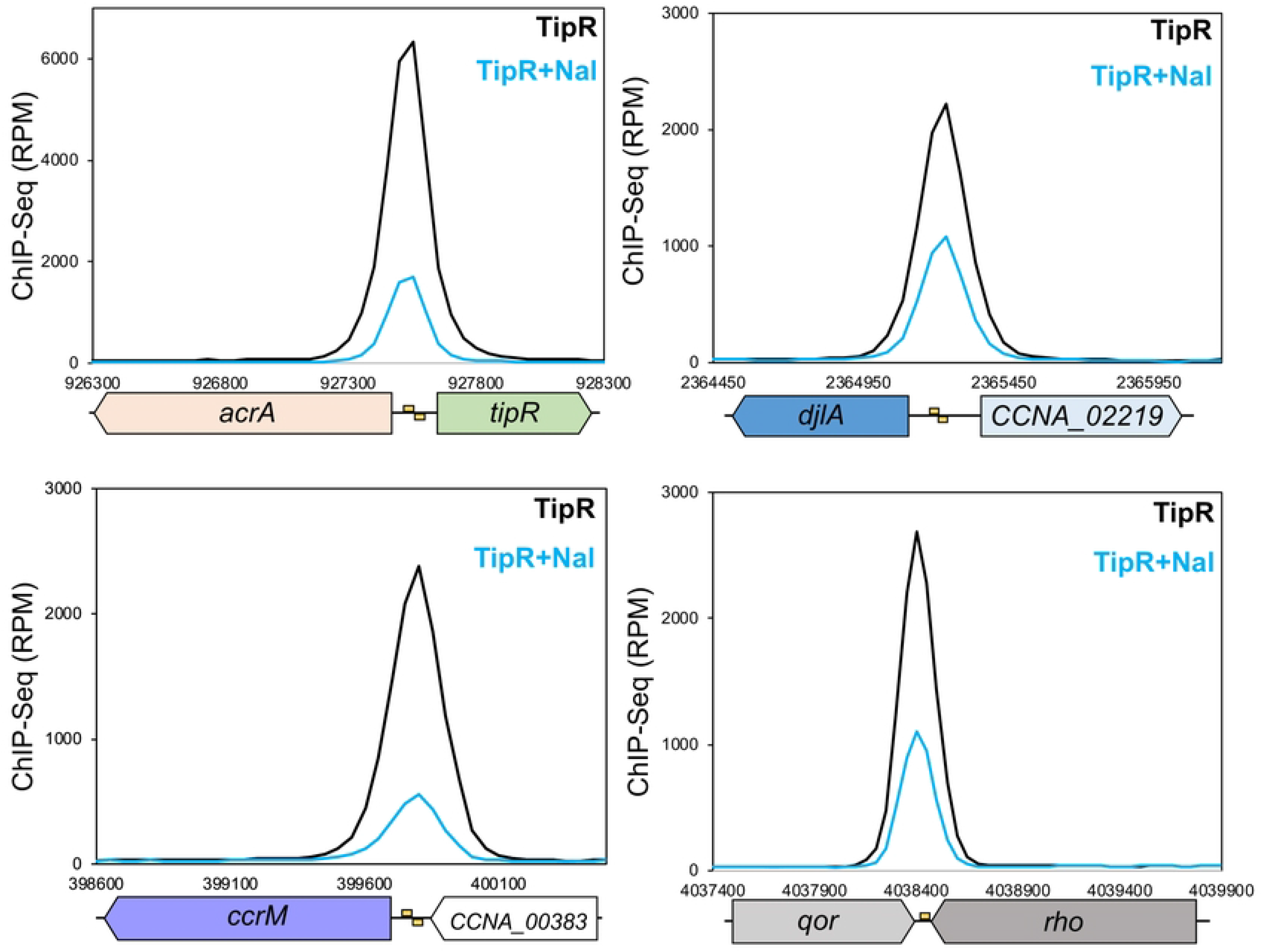
Occupancy of TipR on its chromosomal targets determined by ChIP-Seq. Representation of the reads (in reads per million (RPM)) obtained from the ChIP-Seq analyses covering the TipR binding regions. Positions are indicated under the graphic. Induction was performed for 30 minutes with Nalidixic acid (Nal) 20 µg/mL in PYE (blue line) compared with the not induced condition (black line). The yellow boxes indicate the positions of the putative TipR binding site.

**Figure S7:**
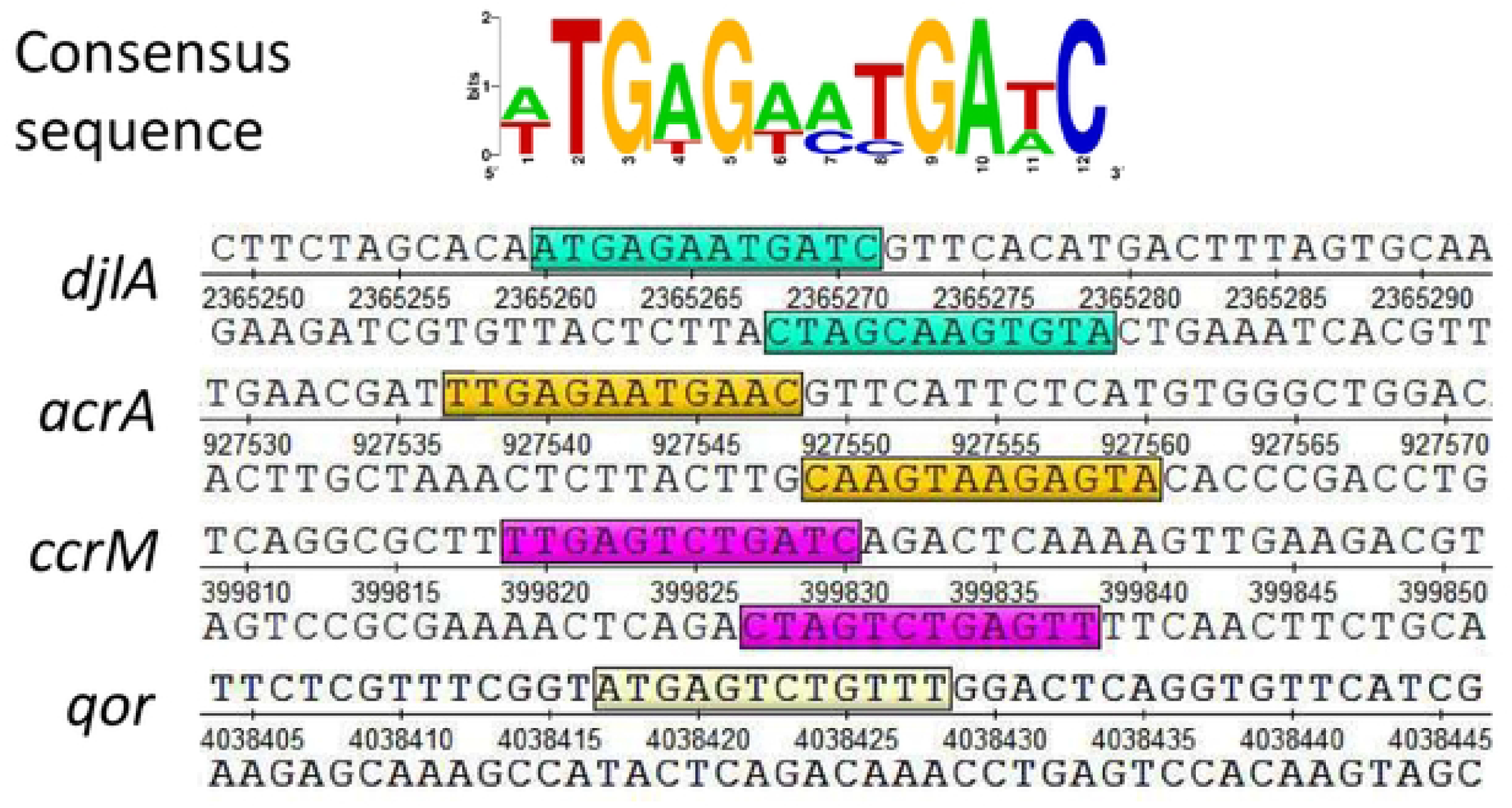
Weblogo analysis of the TipR target DNA sequence. An inverted repeat (IR) sequences was detected at each of the three TipR binding position on the chromosome, plus a half-site at the 4^th^ target site. Consensus sequence of the IRs based on the sequences detected at position 399827, 927547 and 2365267 on the chromosome. This consensus has been identified using MEME software (mean P-value: 4.48×10^-6^) and drawn by WebLogo (crooks).

**Figure S8:**
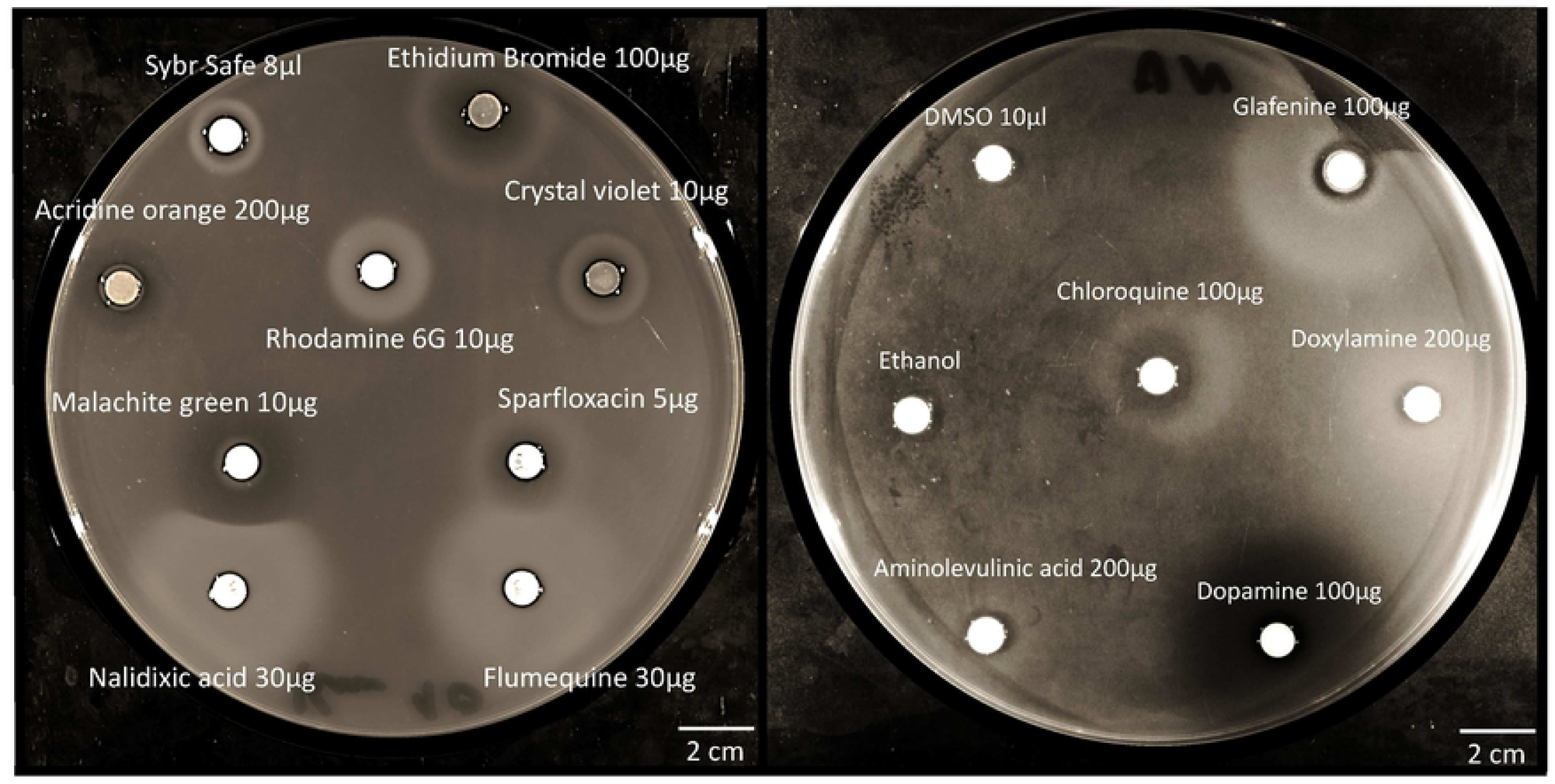
Screen for chemical inducers of P_*acrA*_ using the P_*acrA*_-*nptII* reporter. Reporter assay of P_*acrA*_ activity fused with the *nptII* gene (P_*acrA*_-*nptII* conferring kanamycin resistance) cloned on plasmid plac290 (pP_*acrA*_-*nptII*). To identify inducers, chemicals were spotted on *WT* cells carrying the pP_*acrA*_-*nptII* reporter plasmid embedded on soft agar on PYE plates both containing with kanamycin 10 µg/mL. Plates were incubated for 2 days at 30°C.

**Figure S9:**
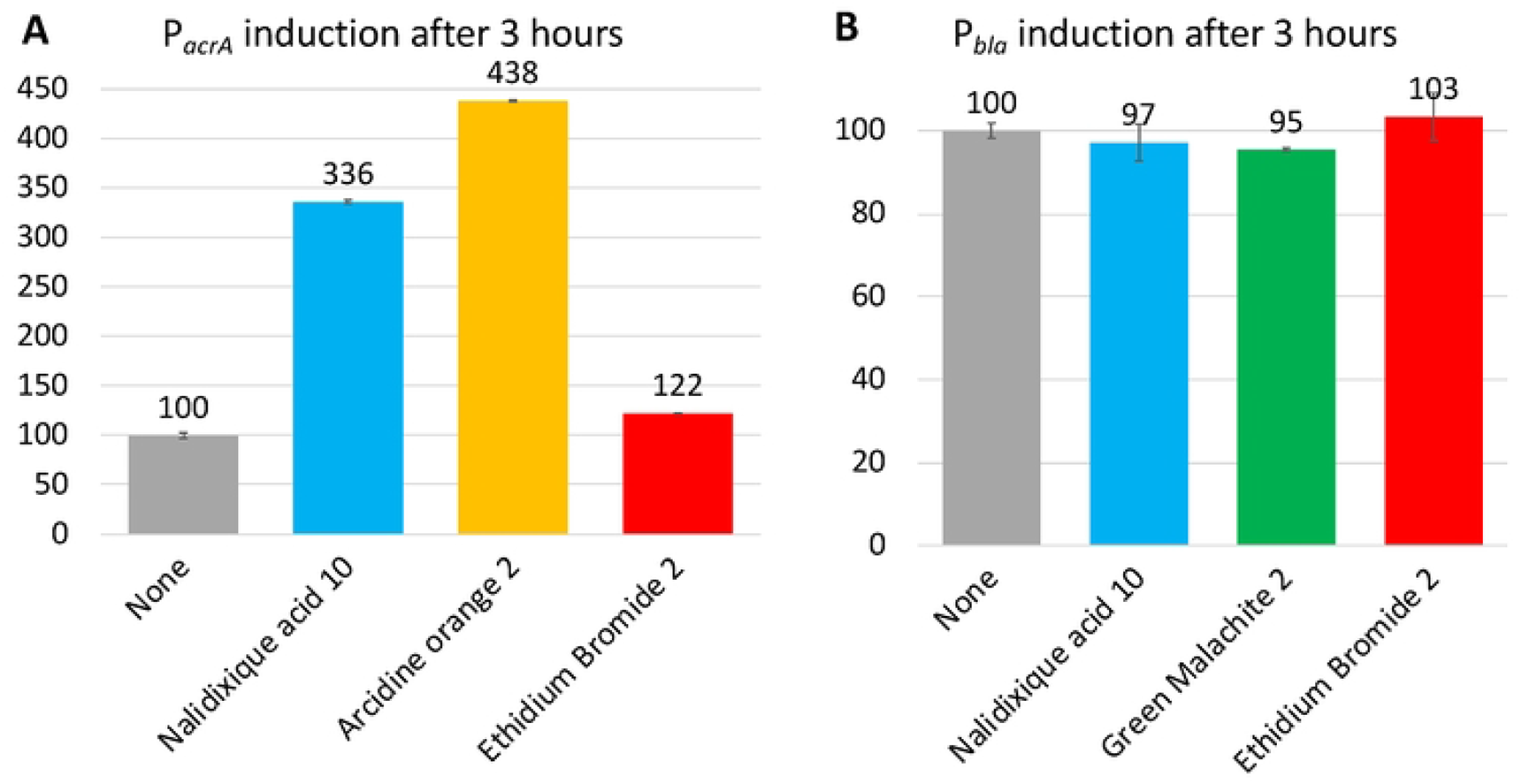
Quantification of P_*acrA*_-*lacZ* activity by different chemical inducers. A) β-galactosidase activity using the P_*acrA*_-*lacZ* in the Δ*acrAB-nodT* cells (A) or P*_bla_-lacZ* in *WT* cells (B). Inductions were performed for 3 hours. All levels are indicated in percentage of expression regarding the basal level of the uninduced state.

**Figure S10:**
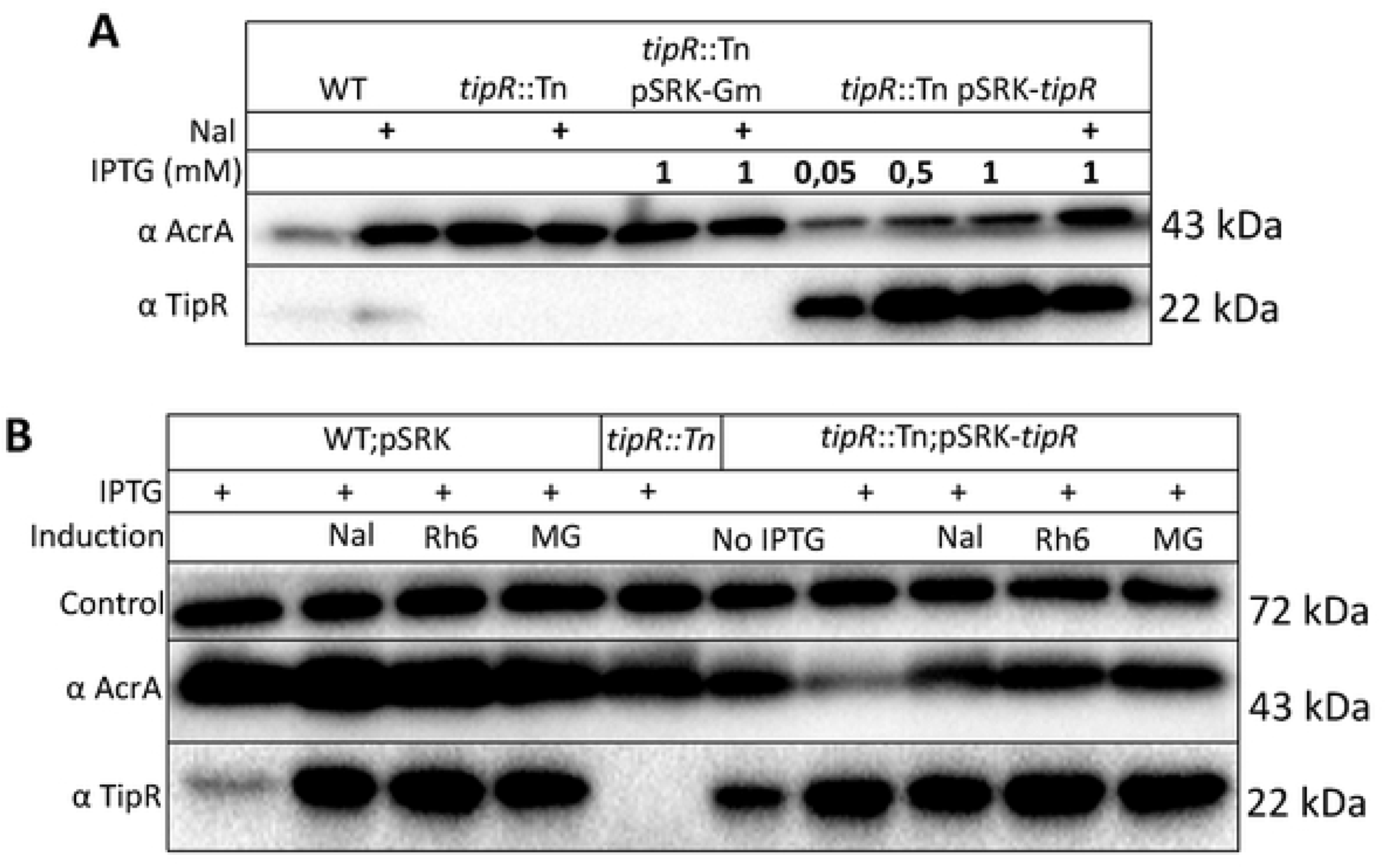
AcrA abundance upon the addition of chemical inducers of P_*acrA*_. A) Immunoblot analysis using anti-AcrA and anti-TipR antibodies to detect AcrA and TipR in *tipR*::Tn cells complemented with the pSRK-*tipR* plasmid, induced with the indicated amount of IPTG (in mM) and Nalidixic acid (Nal, 10 µg/mL) for 2 hours. B) Immunoblot anti-AcrA and anti-TipR against multiple strains grown in PYE with IPTG 0.5 mM, with and without 2 hours induction of Nalidixic acid (Nal, 10 µg/mL), Rhodamine 6G (Rh6, 2 µg/mL) and Malachite Green (MG, 2 µg/mL).

**Figure S11:**
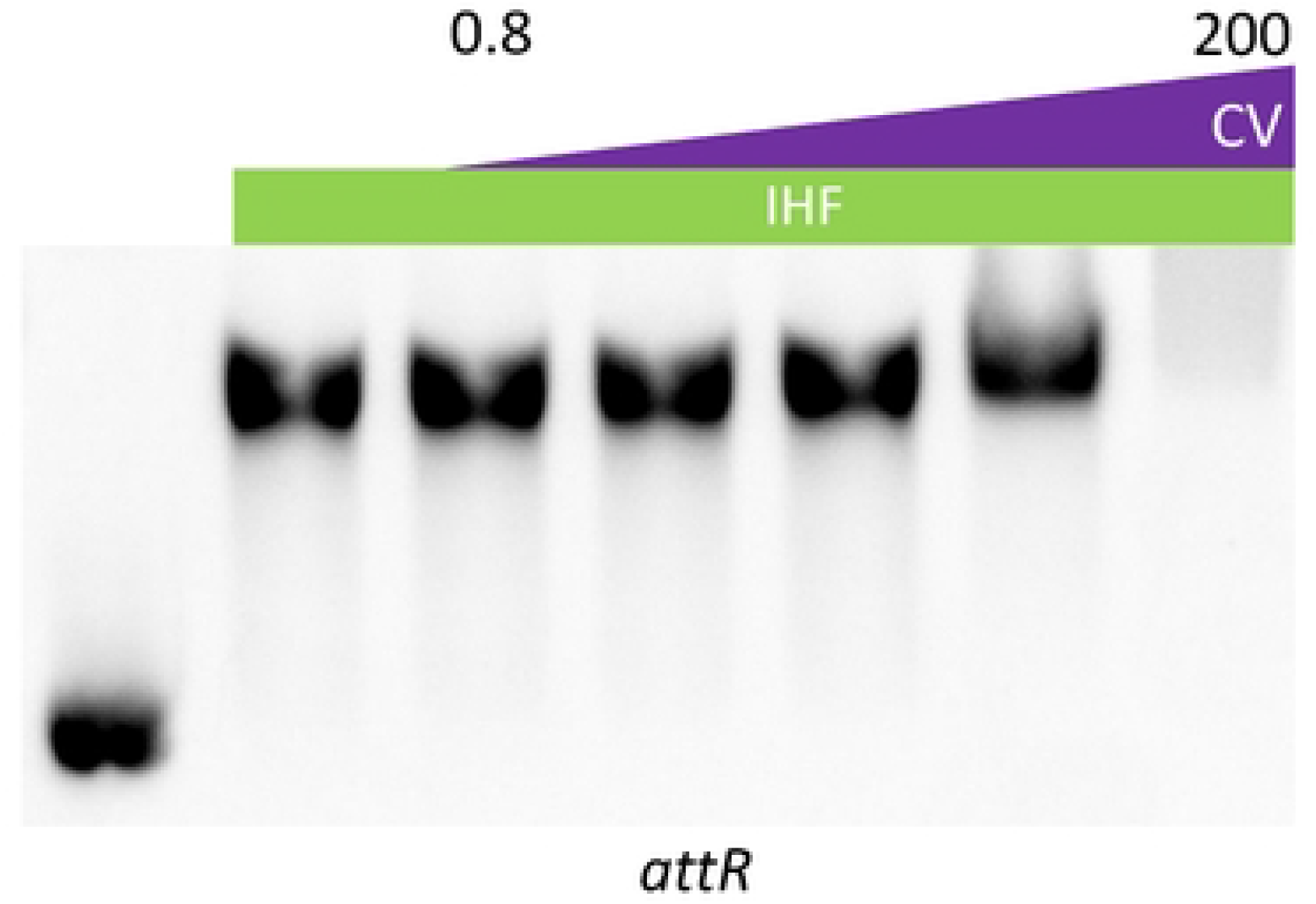
Crystal violet does not affect binding of IHF to its target. EMSA with 4 µM of IHF protein and 200 ng of Cy5-labelled *attR* DNA as probe. All quantities of Crystal Violet (CV) indicated are in µg/mL.

**Figure S12:**
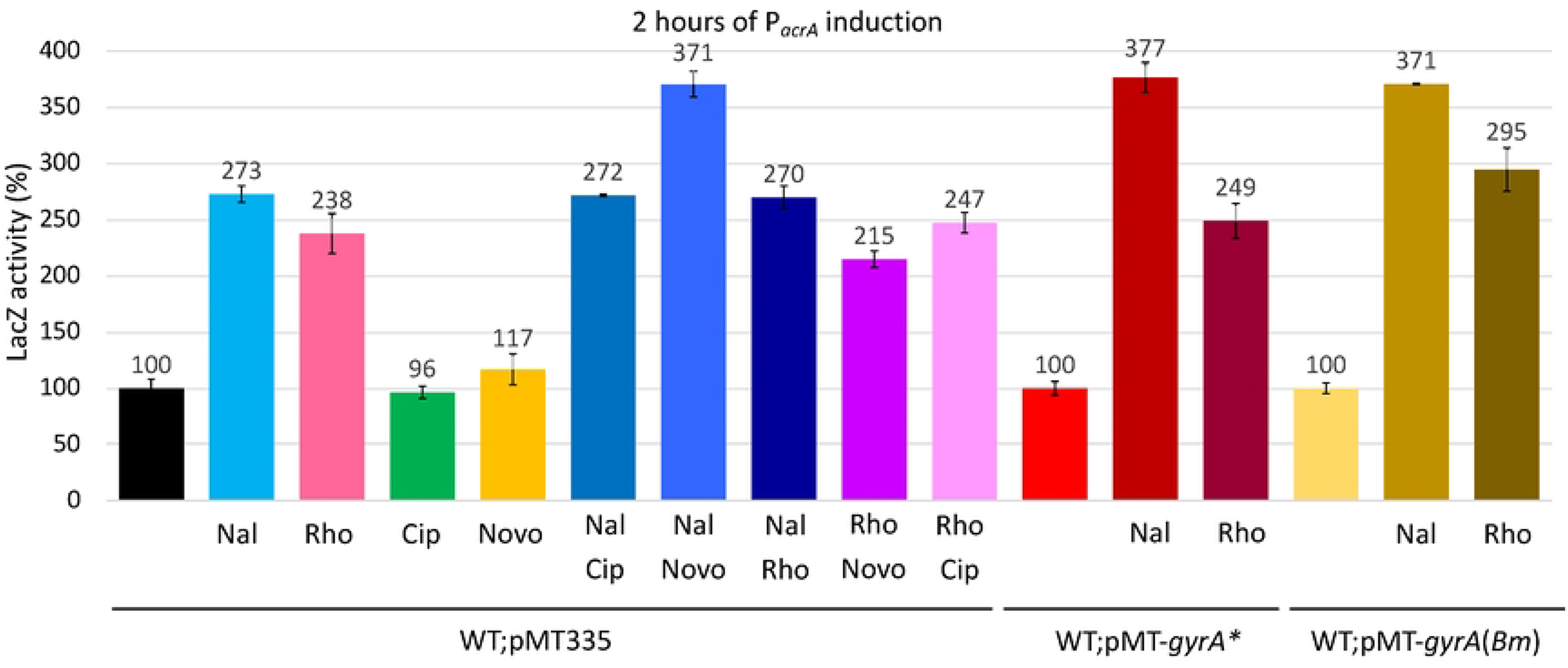
Effect of chemical inducers and *gyrA* variants on P_*acrA*_-*lacZ*. β-galactosidase activity measurements from P_*acrA*_-*lacZ* in NA1000 (*WT*) carrying the pMT335 or a derivativ to express a *gyrA*F96N (GyrA*) or *gyrA* from *Brucella melitensis* (*Bm*) grown in PYE Van 50 µM. Antibiotics were used at the following concentrations: Nalidixic acid (Nal, 10 µg/mL), Ciprofloxacin (Cip, 2 µg/mL), Rhodamine 6G (Rho, 2 µg/mL), Novobiocin (Novo, 10 µg/mL). All levels are indicated in percentage of expression regarding the basal level of *WT* before induction.

**Figure S13:**
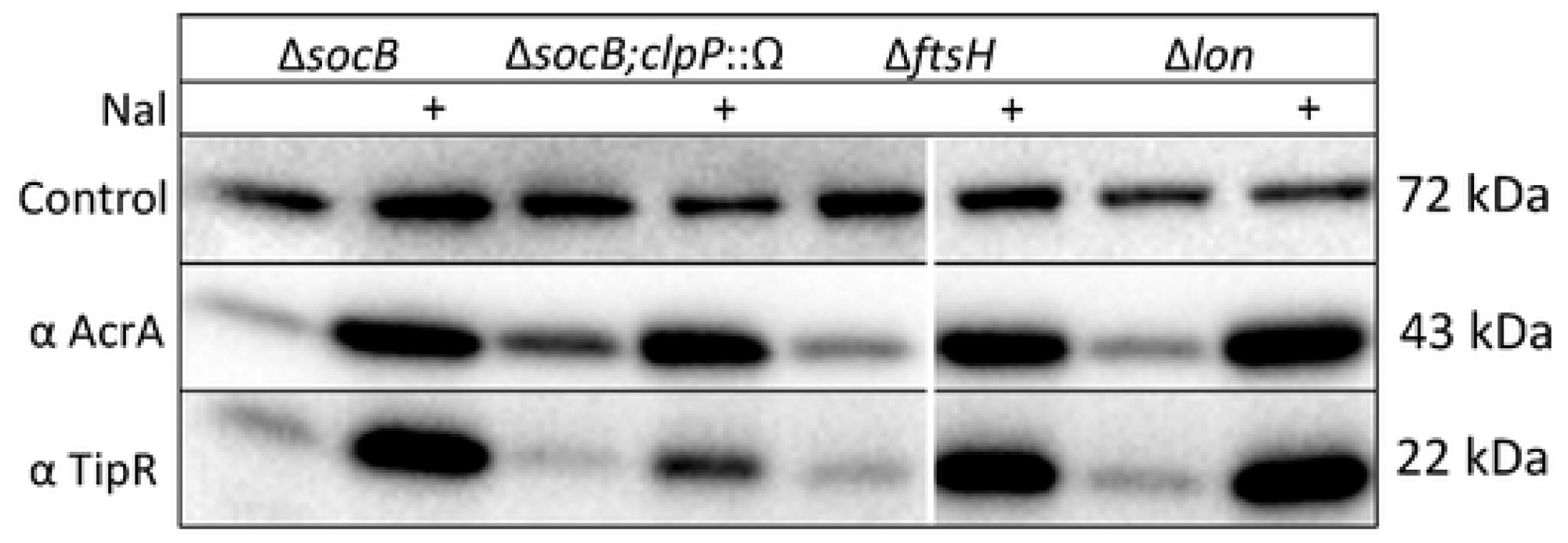
Abundance of AcrA and TipR in protease mutants. Immunoblot analysis using anti-AcrA and anti-TipR antibodies to determine the steady-state levels of AcrA and TipR in various protease mutants of *C. crescentus,* before and after induction with Nalidixic acid (Nal, +, at 10 µg/mL) for 2 hours in PYE. Loading control is anti-CCNA_00164. All samples were loaded on the same immunoblot.

**Figure S14:**
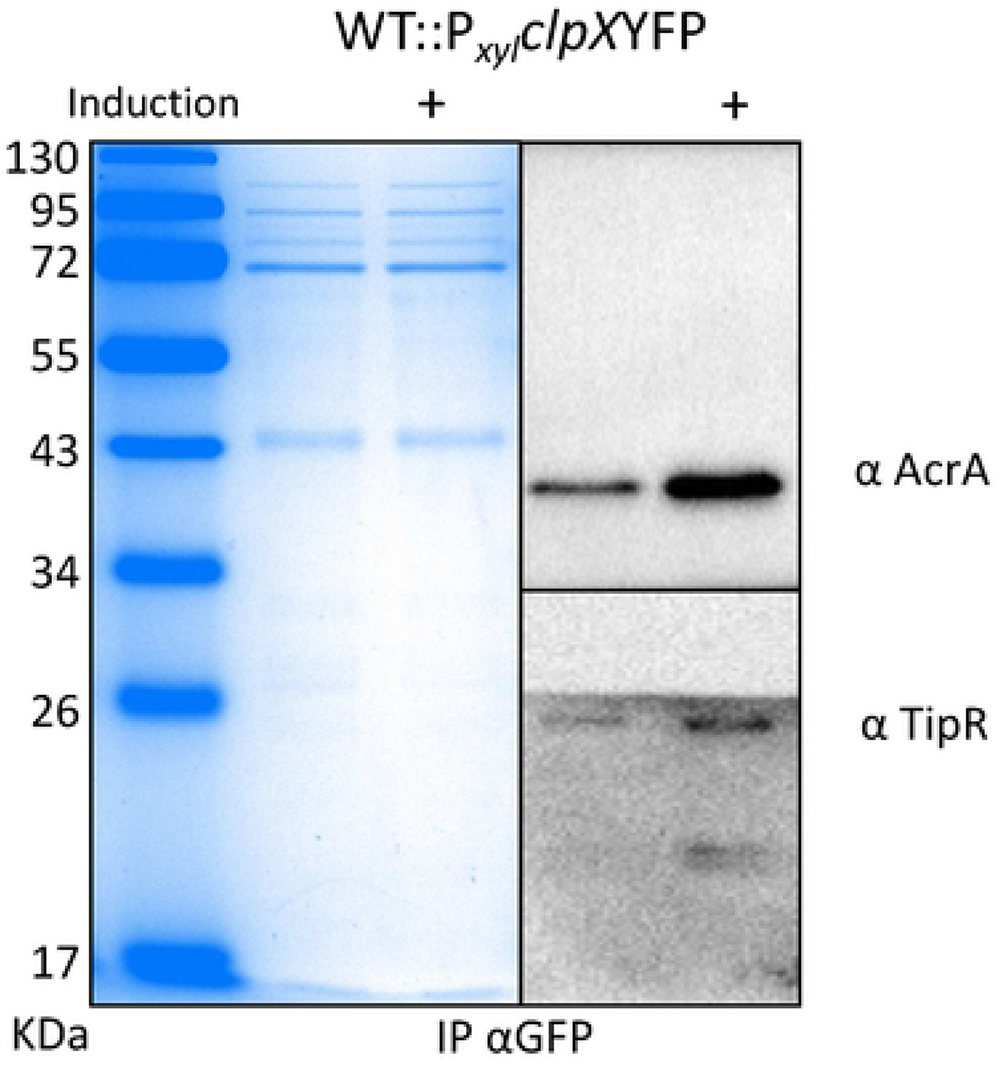
Pull-down of ClpX-YFP. Coomassie Blue stained PAGE (12% gel) (left) and immunoblot (right) of a ClpX-YFP co-immunoprecipitation (GFP Trap Matrix) with anti-TipR and anti-AcrA. Induction with (+) or without (-) nalidixic acid 10 µg/mL.

**Figure S15:**
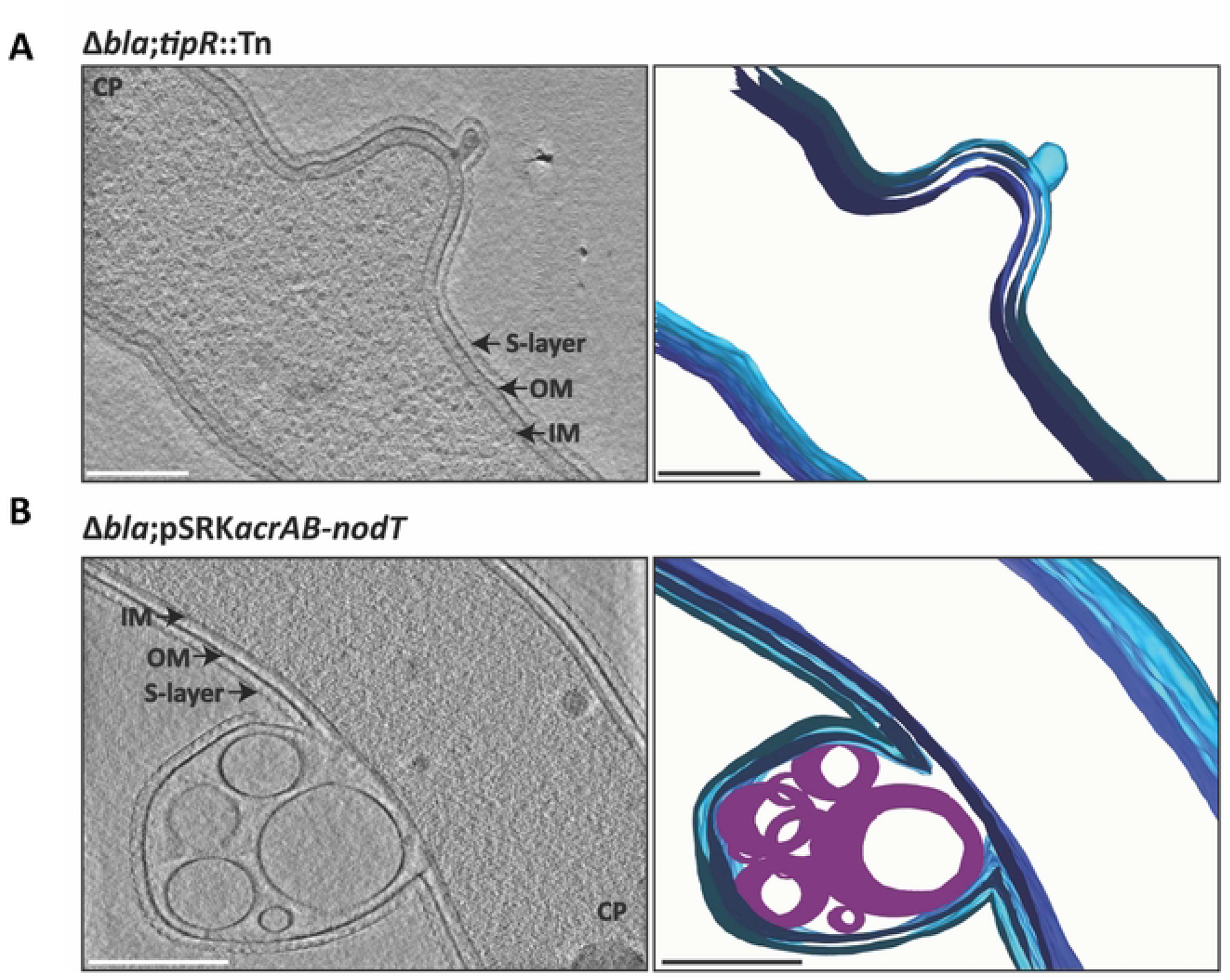
Membrane deformation induced by AcrAB-NodT overexpression. ECT images(left), 3D rendering (center) and superposition of both (right) of *C. crescentus* Δ*bla*;*tipR*::Tn (A) and Δ*bla*;pSRK-*acrAB*-*nodT* (B) cells, imaged by ECT. Scale bar, 250 nm. IM (dark blue), inner membrane; OM (blue), outer membrane; S-Layer (cyan); CP, cytoplasm; Lipid droplet (purple).

**Figure S16:**
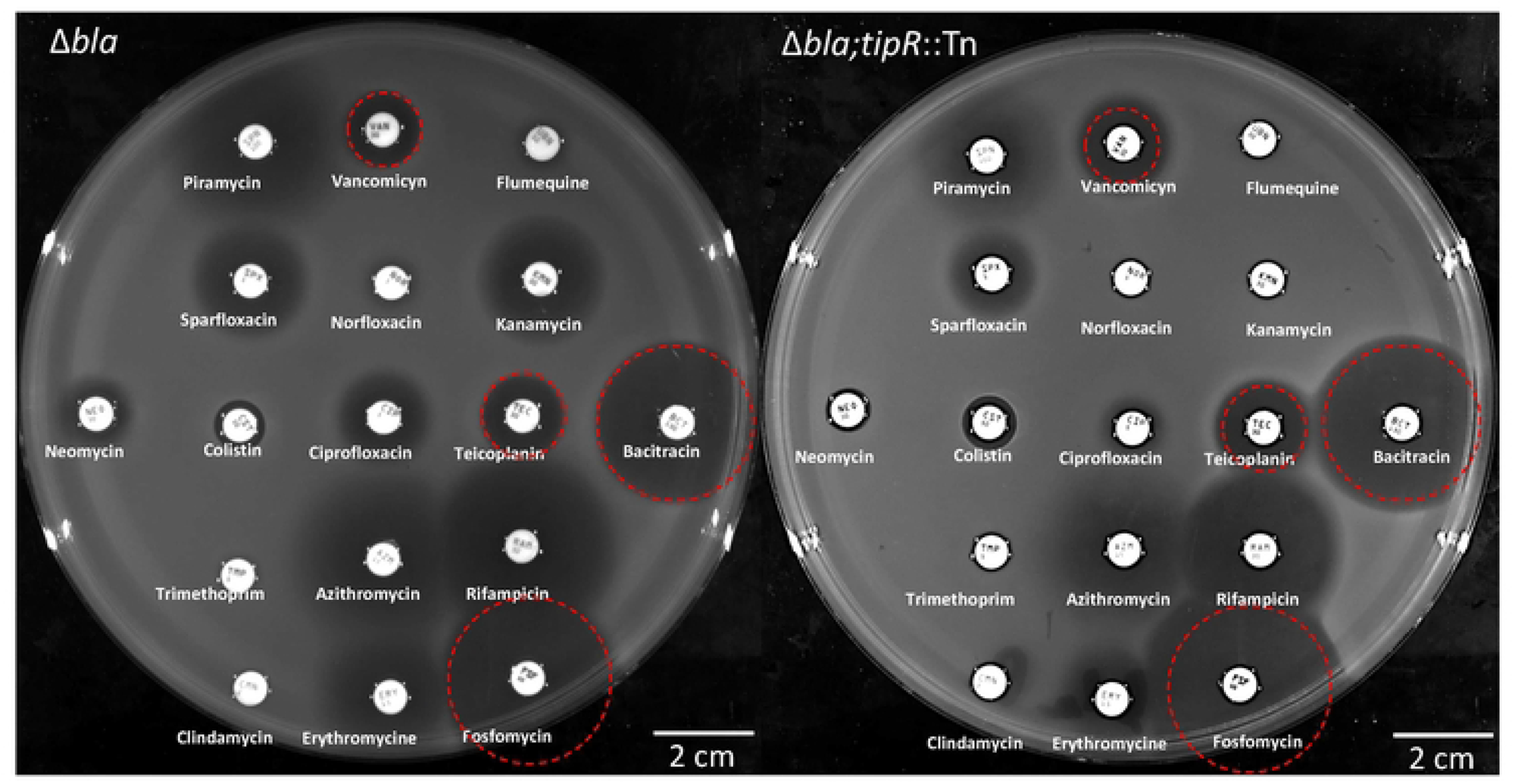
Antibiotic sensitivity of cells overexpressing AcrAB-NodT. Kirby-Bauer based disc diffusion assays to establish antibiograms of *C. crescentus* strains on PYE. Antibiotic discs, from top left to bottom right: Piramycin 100 µg, Vancomycin 30 µg, Flumequine 30 µg, Sparfloxacin 5 µg, Norfloxacin 5 µg, Kanamycin 20 µg, Neomycin 30 µg, Colistin 50 µg, Ciprofloxacin 5 µg, Teicoplanin 30 µg, Bacitracin 130 µg, Trimethoprim 5 µg, Azithromycin 15 µg, Rifampicin 30 µg, Clindamycin 2 µg, Erythromycin 15 µg, Fosfomycin 50 µg. Note: Kanamycin and neomycin resistance is conferred by the *nptII* gene located the transposon inserted in the *tipR* gene. Red circles demarcate the growth boundary for the reference strain.

**Figure S17:**
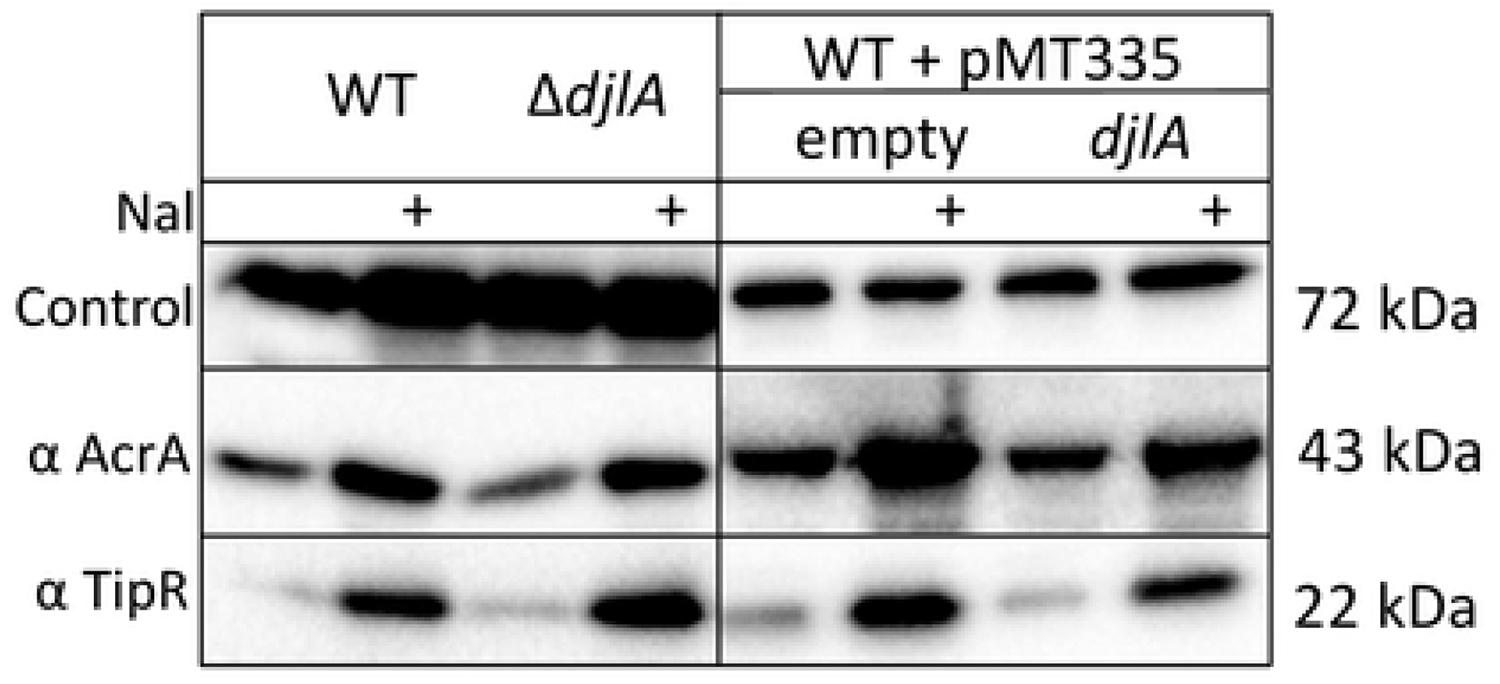
Induction of AcrA and TipR in cells lacking DjlA. Immunoblot analysis of extracts from *WT* and co-chaperone mutants using antibodies TipR and AcrA. All inductions (+) were performed for 2 hours with 10 µg/mL of nalidixic acid (Nal). Strains carrying the pMT335 are induced with vanillate 100µM (Van). Immunoblots performed with antibodies to CCNA_00163 serve as loading controls.

**Figure S18:**
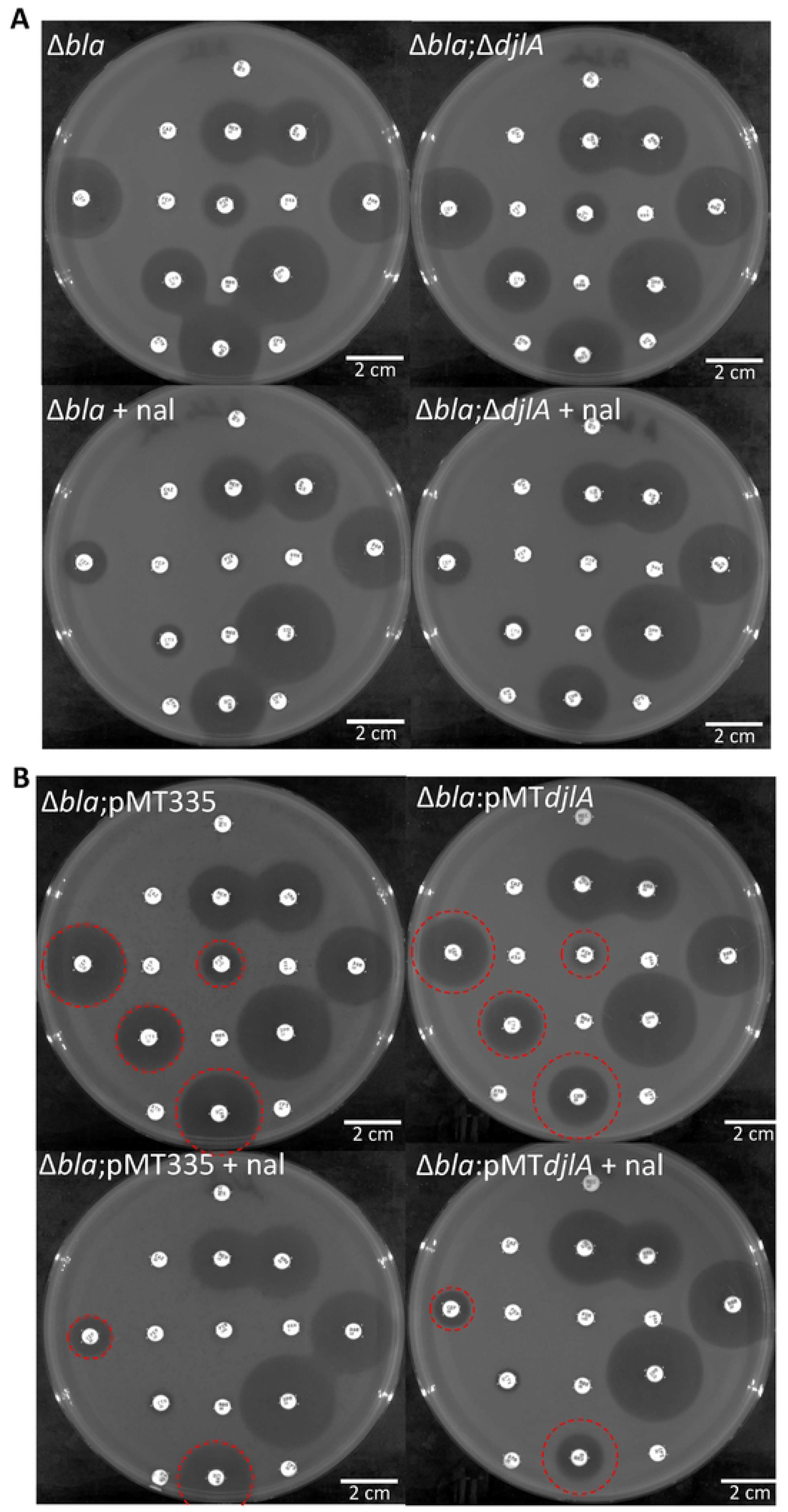
Antibiogram of cells lacking or overexpressing DjlA. Antibiograms of *C. crescentus* strains using antibiotic discs, from top left to bottom right, Mecillinam 10 µg, Ceftazidime 40 µg, Meropenem 10 µg, Amoxicillin 20 µg, Cephalothin 30 µg, Cefepime 30 µg, Piperacillin 100 µg, Oxacillin 5 µg, Doripenem 10 µg, Cefotaxime 30 µg, Moxalactam 30 µg, Imipenem 10 µg, Aztreonam 30 µg, Cephalexin 40 µg, Cefsulodin 30 µg. Nalidixic acid (Nal) induction performed with 10 µg/mL. All plates with strains carrying a pMT335 or pMT335-*djlA* are supplemented with vanillate 100 µM.

**Figure S19:**
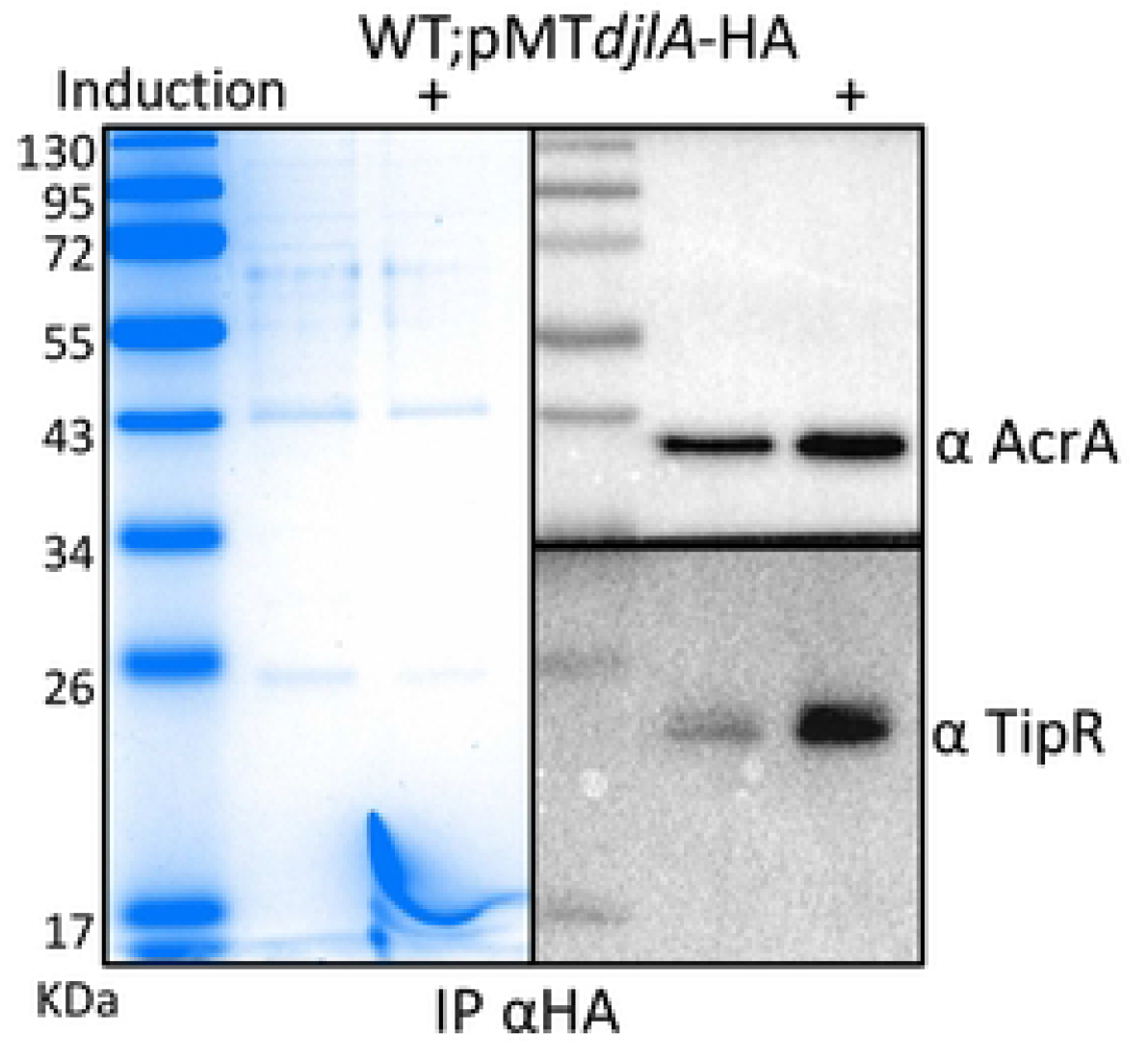
Pull-down of DjlA-HA. Coomassie staining after 12% PAGE (left) and immunoblotting (right) of a DjlA-HA co-immunoprecipitation (using the anti HA affinity matrix) with anti-TipR and anti-AcrA antibodies. Induction was with by nalidixic acid (Nal, +, 10 µg/mL for 2 hours).

**Figure S20:**
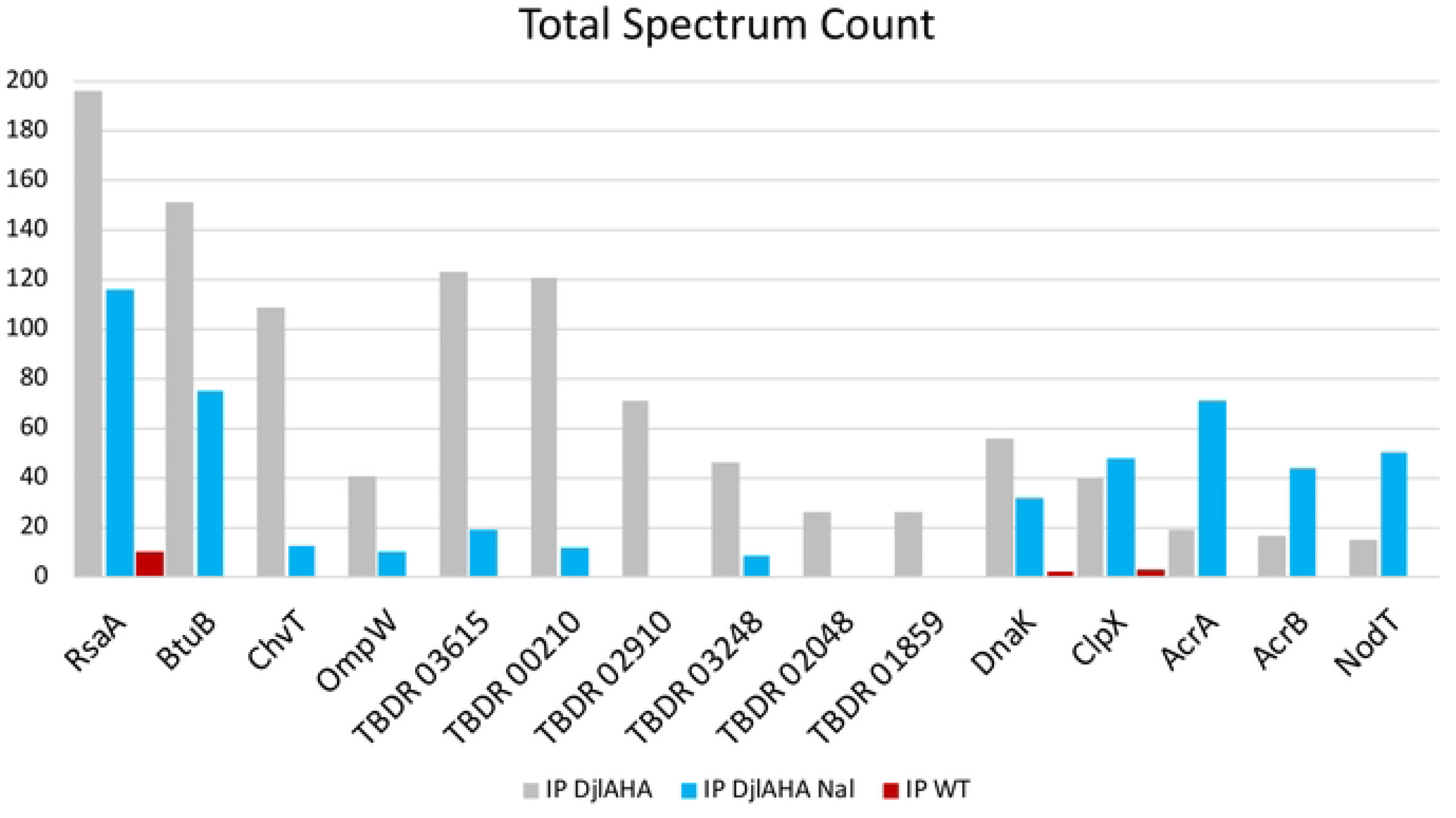
LC-MS/MS analysis of DjlA-HA pull-downs. Graphical representation of the total spectrum count of various proteins detected by LC-MS/MS upon immunoprecipitation of DjlA-HA from soluble cell lysates using anti-HA affinity matrix. The lysates used were from *WT* expressing DjlA-HA plus (Blue) and minus (Grey) NAL, and a *WT* control without DjlA-HA (Red). All the TonB-Dependant Receptor (TBDR) are annotated with their respective CCNA_#.

**Figure S21:** AcrAB-NodT promotes growth of C. crescentus in the cicinity of an unknown fungus. *C. crescentus* strains grown in contact with an unknown fungus on a PYE plate. The fungus was pre-grown the plate for 2 weeks, after which the C. crescentus strains were streaked from the edge of the plate towards the center until in contact with the fungus. The plate was then incubated at 30°C for another 72 hours. The fungus had been isolated in the laboratory as a contaminant on a PYE plate. AcrAB-NodT encoded in the *C. crescentus* chromosome protects against (a) compound(s) produced by the fungus to enable growth of *C. crescentus* in the vicinity of the fungus.

## Notes

### Competing Interest Statement

The authors have declared no competing interest.

